# Sustainability of seed harvesting in wild plant populations: an insight from a global database of matrix population models

**DOI:** 10.1101/2023.01.12.523821

**Authors:** Anna Bucharova, Oliver Bossdorf, J. F. Scheepens, Roberto Salguero-Gómez

## Abstract

Seed harvesting from wild plant populations is key for ecological restoration, but may threaten the persistence of source populations. Consequently, several countries have set guidelines limiting the proportions of harvestable seeds. However, these guidelines are so far inconsistent, and they lack a solid empirical basis. Here, we use high-resolution data from 298 plant species to model the demographic consequences of seed harvesting. We find that the current guidelines do not protect populations of annuals and short-lived perennials, while they are overly restrictive for long-lived plants. We show that the maximum possible fraction of seed production – what can be harvested without compromising the long-term persistence of populations – is strongly related to the generation time of the target species. When harvesting every year, this safe seed fraction ranges from 80% in long-lived species to 2% in most annuals. Less frequent seed harvesting substantially increases the safe seed fraction: In the most vulnerable annual species, it is safe to harvest 5%, 10% or 30% of population seed production when harvesting every two, five or ten years, respectively. Our results provide a quantitative basis for seed harvesting legislations worldwide, based on species’ generation time and harvesting regime.

**Significance:** The UN Decade on Ecosystem Restoration, 2021-2030, foresees upscaling restoration, and the demand for native seed is skyrocketing. Seeds for restoring native vegetation are often harvested in wild, but too intensive harvest can threaten the donor populations. Existing guidelines that set limits to wild seed harvest are mostly based on expert opinions, yet they commonly lack empirical basis and vary among regions in one order of magnitude. We show that the current guidelines urgently need to be reformulated, because they are overly restrictive in long-lived species, while they do not protect annual plants from extinction. Using matrix population models of nearly 300 plant species, we provide a quantitative basis for a new seed harvesting legislation world-wide.

## Introduction

The restoration of degraded ecosystems is a major goal of global nature conservation (1). We are in the middle of the ’UN Decade on Ecosystem Restoration’ (2), with a key goal to reverse the destruction and degradation of billions of hectares of ecosystems. However, ecological restoration at such scales requires high volumes of plant seeds for the re-establishment of native vegetation (3).

Although there is a growing industry for the production of wild plant seeds in specialised seed orchards (4, 5), large-scale harvesting of seeds from wild populations is still common in ecological restoration, and is projected to continue growing (6). Seed harvesting is particularly common for plant species that are long-lived or difficult to cultivate (7–10).

With increasing demands for wild plant seeds, there is a growing risk of driving source populations to local extinction (11, 12). Moreover, donor populations are often remnants of habitats with high conservation value (11, 13). Some regions, in particular the US (14), Australia (15), and Europe (16, 17), have therefore begun to set limits for the maximum fraction of seeds that can be harvested annually from wild plant populations, to prevent significant negative effects on their long-term viability (‘*safe seed fraction*’, hereafter). Notably, though, the safe seed fraction guidelines are inconsistent across countries, with *e.g.* 20% harvest allowed in the US (14) and 10% in Australia (15), but only 2-10% in Germany, depending on plant growth type (16). When the harvest does not take place annually, some guidelines permit higher safe seed fractions (16). In general, however, these guidelines are mostly based on expert opinion and lack a solid quantitative basis.

Only a few studies have experimentally tested the effects of seed harvesting on wild populations. However, these studies are either focused on individual species or specific ecosystems (11, 18, 19). More effective rules would require collection of data across multiple species and ecosystems, but this course of action is labour- and cost-demanding. As an alternative to collecting new data, Menges and colleagues (12) used published plant matrix population models to link seed harvesting to the probabilities of population extinction for 22 perennial species. While this study is widely used to back up seed collection guidelines for rare species for ex-situ conservation (e.g., (15, 20), the species set is limited to mostly herbaceous perennials of temperate and subtropical North America. To obtain a quantitative basis for predicting the effects seed harvesting on wild populations globally, data from many more species across life histories and ecosystems are essential.

Here, we employed a modelling approach and simulated seed harvesting for 298 plant species ranging from annuals to long-lived trees from many habitats around the globe using matrix population models stored in the the COMPADRE Plant Matrix Database (21, 22), Table S1.

Specifically, we (1) tested the efficacy of current guidelines at safeguarding long-term population persistence, (2) identified traits that are associated with species vulnerability to seed harvesting, and (3) used the trait that best determines species vulnerability to seed harvesting, generation time, to predict safe seed fraction, and formulated quantitative basis for seed harvesting in wild plant populations world-wide.

## Results

To test how well the current safe seed fraction guidelines protect source populations from overharvesting, we modelled the maximal possible harvest fractions permitted in the US, Australia, and Germany. To allow comparison across species, we expressed effects of seed harvesting as relative population sizes, where 1 indicates no effect, 0 indicates extinction, and *e.g.* 0.8 represents a 20% reduction of population size in comparison to the population size that would be reached without seed harvesting. Seed harvesting according the existing safe seed fraction guidelines results in rather variable relative population sizes among species (Figure 1). For instance, the current US guidelines (20% seed harvesting allowed) protect long-lived palms, with relative population sizes of 0.6 to 1 after 30 years, but would drive all 10 annual plants in our data to extinction (Fig. 1). With the more restrictive German guidelines (2% seed harvesting allowed), annual plants are projected to persist, with relative population sizes of 0.54 to 0.63 after 30 years. Within all other plant growth types, the effects of seed harvesting on the relative population sizes are much more variable. For example, with the 20% seed harvesting currently allowed in the US, the predicted relative population sizes of herbaceous perennials would range from 0 (local extinction) to 1 (no effect) after 30 years, while that of shrubs would range from 0.12 to 0.99, of succulents from 0.27 to 0.99, and of trees from 0.18 to 0.99 (Fig. 1).

**Figure 1.**
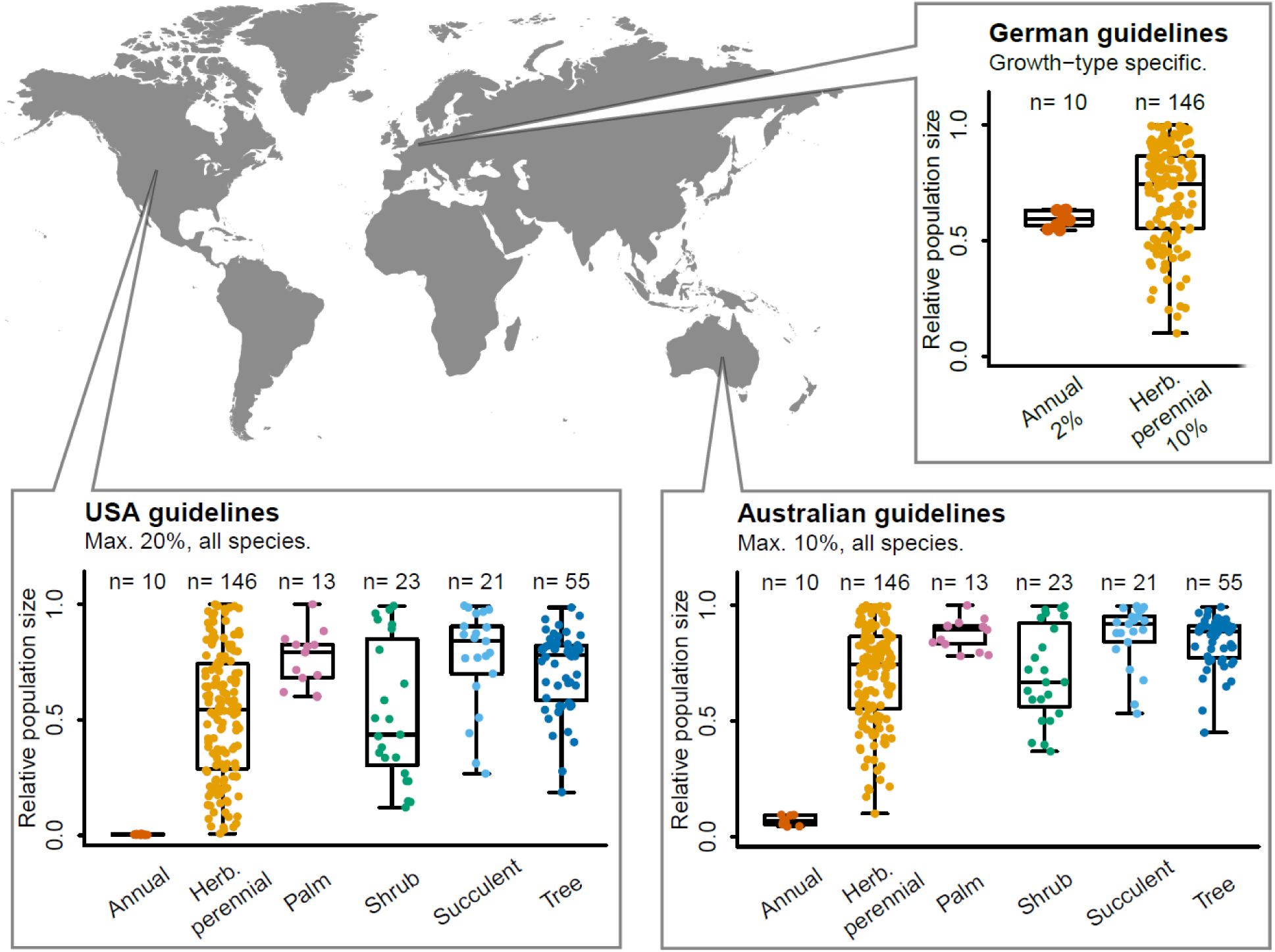
Predicted effects of 30 years of continuous seed harvesting on the relative population sizes of 298 plant species worldwide, using the current guidelines of countries where legislation exists: USA, Germany, and Australia. Points represent individual species. The data result from simulation of seed harvesting using matrix population models parameterised with data from natural populations. Herb. = Herbaceous; n = numbers of species included.

We next examined whether and which life history traits are better predictors of seed harvesting impacts (Figure 2). We found out that generation time, the mean age of reproductive individuals in the population, is the strongest predictor of population vulnerability to seed harvesting. This life history trait alone explains 52.3% of the variation in harvesting vulnerability, and vulnerability to seed harvesting decreases with increasing generation time (Fig. 2B). Four other life history traits are also significantly related to seed harvesting vulnerability (Fig. 2B) – species that reproduce more frequently and/or postpone their first reproductive event are more vulnerable to seed harvesting, while species with clonal reproduction and/or persistent seed banks are less vulnerable – but the predictive power of these traits is low (Fig. 2A, Table S3). Population vulnerability also differs significantly among plant growth types, but with minor effects (Fig. 2C, Table S3). All five life history traits together explain 62.3% variability in vulnerability to seed harvesting among species.

**Figure 2.**
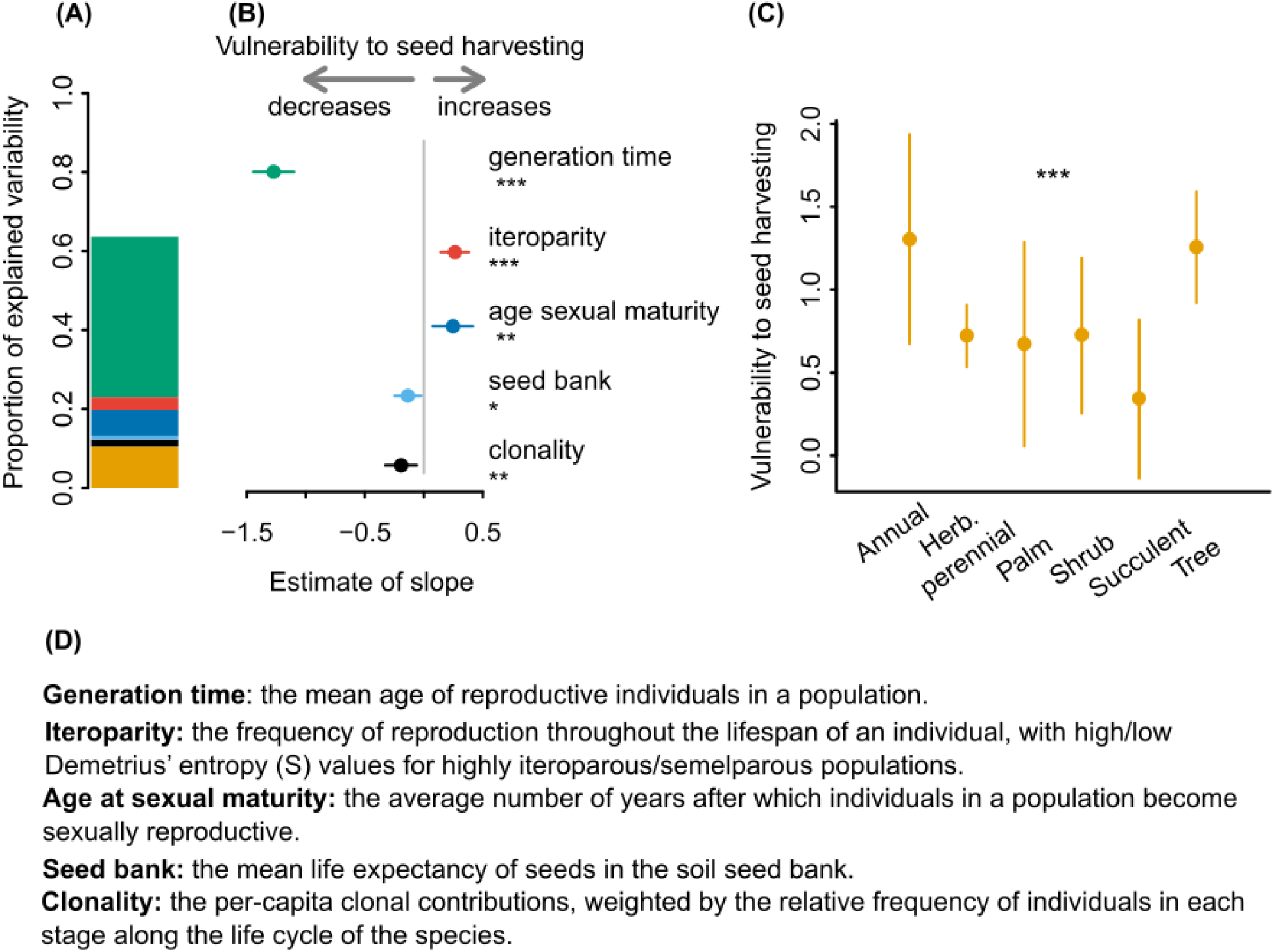
Associations of plant life histories and growth forms with variation in seed harvesting vulnerability across 298 plant species, as calculated from matrix population models parameterised with data from natural populations. (A) Proportion of variability explained by different life history traits, and (B) their effect estimates. (C) The fitted values of vulnerability for different growth types. Estimates in (B) and (C) are presented with their 95% credible intervals. As both vulnerability to seed harvesting and all explanatory variables were standardised prior the analysis, the slope estimates are in arbitrary units. (D) Definitions of the five examined life history traits (for calculation see Table S2). Herb. = herbaceous. Significance levels: * P<0.05, ** P<0.01, *** P<0.001, See Table S3 for detailed model results.

To improve the efficacy of seed harvesting regulation, we then used the best predictor of species vulnerability to seed harvesting, generation time, to estimate safe seed fraction across species. For annual harvesting, the safe seed fraction ranges from close to 0% to 100%, with an average of 2.3% (95% CI: 0.5-4.1%) for annual and biennial plants, 10.1% (6.8-14.2%) for species with a 5-year generation time, and 40.1% (36.4-43.7%) for species with generation times of 20 years (Fig. 3A). With simulated harvesting only every two years, the safe seed fraction for annuals and biennials increases from 2.3% to 5.3 % (2.7-7.9%), and with a 5-year or 10-year harvesting interval to 11.3% (6.5-16.0%) and 30.3% (23.8-36.8%), respectively (Fig 3B-D). For plant species with generation times above two years, a 5-year harvesting cycle resulted in an average safe seed fraction of >30% (Fig. 3C). While safe seed fraction critically depends on generation time, there is substantial residual variation among species.

**Figure 3.**
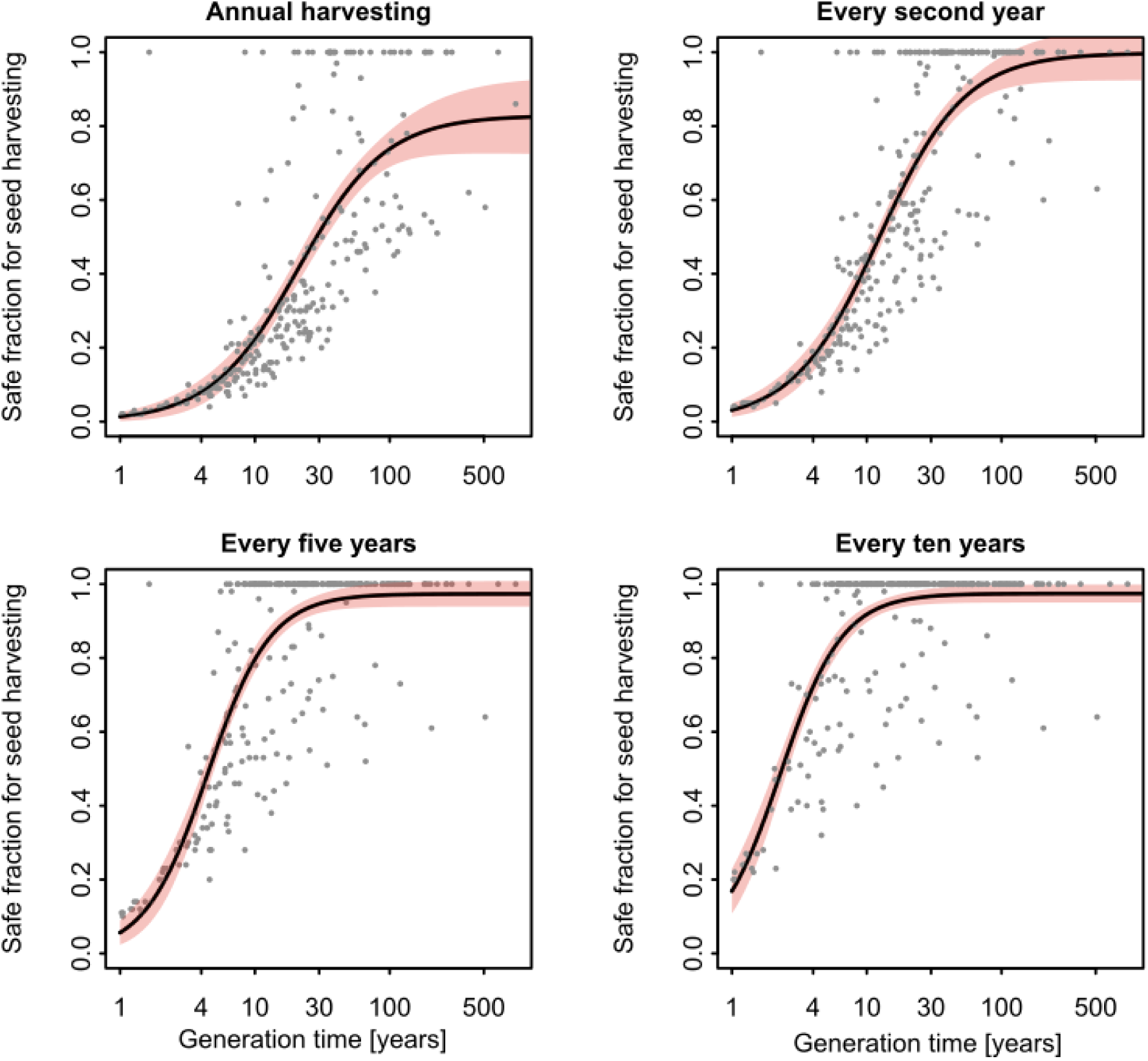
Relationships between the generation times of 298 plant species and their safe fractions for seed harvesting, estimated at different harvesting frequencies. The safe seed fraction is the maximum proportion of annual seed production of a population that can be harvested without reducing the relative population size to below 50% in 30 years.

The estimated safe seed fraction for each species was not substantially affected by environmental stochasticity. The median of safe seed fractions based on models that included environmental stochasticity (see methods) was on average 1.8% larger than the safe seed fraction based on the mean models for each species, yet they were closely correlated (Figure S3).

## Discussion

Seed harvesting in wild population is indispensable for ex-situ conservation and ecosystem restoration, but overharvesting can threaten source populations (13). Consequently, some countries have introduced limits that restrict wild seed harvesting (14–16). Here, using data from wild populations of 298 plant species from five continents, we show that the current seed harvesting guidelines are often ineffective: existing guidelines do not protect populations of annuals and short-lived perennials, while they are overly restrictive for long-lived plants. Based on generation time, the trait that best predicts seed harvesting vulnerability, we estimate that safe seed fraction varies from 2% in annual and biennial plants to 80-100% in long-lived plants, when seeds are harvested annually. Lower frequency of harvesting allows for higher seed fractions in a viable way. The safe seed fractions presented here can serve as a solid quantitative basis for seed harvesting regulations globally.

When wild seed harvesting follows the existing safe seed fraction guidelines, the effects on population sizes can vary from no effect to extinction, depending on the species. For example, annual seed harvesting of 20% of the annual seed production, as currently recommended in the US (14), would have small effect on palms, trees or some herbaceous perennials, but it would drive all annuals plants to extinction within three decades. In reality, extinction will be less common because we modelled an extreme scenario when seeds are harvested every growing season for 30 consecutive years from the same population, which is possible but uncommon (13). Nevertheless, the high variability in model outcomes highlights that effective safe seed fraction guidelines must be more nuanced than one-size-fits-all – one safe seed fraction for all species – as currently implemented in many regions (14, 15, 17).

The current German safe seed fractions guidelines are plant growth-type specific (16). For annual plant species, the safe seed fraction is 2% when harvesting annually, which in our modelling does not cause unacceptable population declines (Figure 1). For herbaceous perennials, the safe seed fraction in Germany is set to 10% for annual harvest, yet this threshold leads to a wide range of relative population sizes, from substantial population declines to no effects. The variability within the herbaceous perennials is even stronger when following the US guidelines (20% of annual seed production). Plant growth type alone is thus a poor predictor of species vulnerability to seed harvesting.

Over 60% of the vulnerability to seed harvesting is predicted by life history traits. The highest predictive value in our analyses offers generation time, which alone predicts the seed harvesting vulnerability by more than 50%. Population growth rates in long-lived species are generally insensitive to changes in fecundity (23, 24). Indeed, (12) showed that long-lived plants are relatively insensitive to seed harvesting. Other life history traits in the present study have much smaller predictive power for seed harvesting impacts. For instance, species with higher iteroparity (*i.e.* reproducing more than once during their life cycle), and species that are later sexually mature, are more vulnerable to seed harvesting, while clonal species and species with permanent soil seed banks are less vulnerable. The buffering effect of seed bank against the effects of seed harvesting are well supported by the literature (25). However, the relatively small effect of clonality on the impacts of seed harvesting is surprising, since clonality provides an alternative reproduction independent of seed production, and has been experimentally identified as a major predictor of vulnerability to seed harvesting in grassland plants (18). This discrepancy is likely because many matrix population models calculate generation times of individuals originated from seeds, *i.e.* genets. Clonal reproduction thus leads to longer generation times of the genets (24, 26), and explains little additional variability in vulnerability to seed harvesting above what is already explained by generation time as the more universal predictor.

To provide a universal quantitative basis for seed harvesting guidelines, we estimated safe seed fraction as a function of generation time, the best predictor of vulnerability to seed harvesting (Figure 3). The lowest safe seed fractions are in annuals and biennial, 2.3% for annual harvest, which is close to the current German guidelines of 2%, (16). The safe seed fraction continuously increases with generation time, but remains below 10% for plants with generation times of five years and less. Adhering to such low seed safe fractions is possible only when collecting seed manually, yet this is very labor intensive. In grasslands, seeds are commonly harvested using combine harvesters, which typically removes 30% of the ripe seeds (27). Such a high proportion is safe for annual harvesting only in species with generation time above 15 years. Grasslands are dominated by annuals and herbaceous perennials, of which 60% in our dataset have generation times below 15 years. Annual seed removal with combine harvesters thus threatens a substantial proportion of grasslands species, especially non-clonal forbs and annuals (18).

Less frequent harvesting allows higher safe seed fractions. Harvesting seeds less often is already suggested as a precautional principle in some guidelines (*e.g.* (13, 17), although mostly without a clear specification of safe seed fractions and harvesting frequencies. Less frequent harvesting is relevant especially for species with short generation times, where the safe seed fraction is the lowest. In annual and biennials, the safe seed fraction increases from 2.3% for annual harvesting to 5% when harvesting every second year, 11% every five years and 30% every 10 years. Importantly, harvesting at 10-year intervals allows to collect 30 % of the seed production even in the most vulnerable species. Seed harvesting with combine harvesters, which collects on average 30% of the seed (27), should be sustainable even in drylands with high proportion of annual plants, if done at sufficiently long intervals.

Seed harvesting is less problematic in species with long generation times. In species with generation times above 20 years (most trees and palms, many shrubs and some herbaceous perennials (28)), safe seed fractions are above 40% when harvesting every year, and above 80% when harvesting less frequently. Previous empirical and modelling studies also reported that long-lived species are rather insensitive to seed harvesting (12, 18, 29), although too frequent and too intense harvesting can deplete populations of seedlings (19). Even in long-lived species, it might thus be beneficial to omit seed harvesting in some years to give populations opportunities for juvenile recruitment.

Our results demonstrate the demographic impact of seed harvesting, and how it depends on plant life histories. Yet, we could have overestimated harvesting impacts for three reasons. First, our analyses are based on matrix population models of species averaged across years and sites, but temporal or spatial variation in demographic rates could buffer some impacts of seed harvesting (30). Indeed, incorporating environmental and demographic stochasticity into our models in a subset of species resulted in safe seed fractions on average 1.8% larger, confirming that matrix averaging may cause overestimation, but the effect was small. Second, our approach assumes plant populations to be seed-limited. However, longer-lived plants are often limited by safe sites rather than seeds, whereas seed limitation is more common in short-lived species (31). It is thus likely that in longer-lived species the effects of seed harvesting are even less severe than our findings suggest, but for annuals and short-lived forbs – the most vulnerable to seed harvesting – our results are more likely to be accurate. A specific case of safe-site limited habitats are European seminatural meadows that are annually mown with the biomass, including a large proportion of seed, used as fodder for domestic animals. Species growing in this ecosystem are likely adapted to regular seed removal and thus less vulnerable to seed harvesting than predicted by our models. Third, our models do not incorporate maximal carrying capacities, because this information is rarely available for matrix population models. In populations with high population growth rates and close to carrying capacity of the environment, matrix models still predict population growth even though the population already reached maximal space occupancy. In such cases, seed harvesting might have much smaller effect than predicted.

Seed harvesting in wild populations should be generally accompanied by monitoring of the harvested sites. Our results provide the currently best quantitative basis for sustainable seed harvesting in wild populations. Yet, they are model results, and all models are simplifications of the reality as it is impossible to capture the full complexity of the real world (32). As a precaution, and to be able to adjust harvesting practice if necessary, it is therefore important to monitor the harvested sites. The safe seed fractions presented here cause only very slow population declines, maximum 2% per year, and monitoring every few years should be sufficient to detect unexpected negative effects on population sizes before the population would be irreversibly damaged.

In summary, we show that seed harvesting in wild populations is possible and allows long-term population persistence, but the harvesting must be guided by the critical factors of plant generation time and harvesting frequency. For longer-lived species, harvesting large fractions of seeds is unlikely to harm wild populations, particularly if seeds are not harvested every year. For short-lived species, though, more caution is necessary. A profitable harvesting of 30% of the seeds of annual species may only be possible if the harvesting takes place only every 10 or more years. However, ultimately, even with improved guidelines, seed harvesting from wild populations is unlikely to cover the growing worldwide needs of ecological restoration (33). The ambitious targets of the UN Decade on Ecosystem Restoration (2) may only be reached with professional, large-scale seed production in seed orchards (4, 34, 35).

## Methods

We used data stored in The COMADRE Plant Matrix Database (version 5.0.0. last accessed 25.8.2019 (22), and selected matrix population models for 298 species (SM, section 1). As the ultimate goal of this study was to simulate seed harvesting, we selected field-based models for angiosperms with clearly defined sexual reproduction (SM 1.3 for details). For the majority of studies in COMPADRE, matrix population models are available for several annual transitions and populations. For all calculations, except the stochastic simulations (see below), we used a single MPM per species averaged across all years and populations available for that species. Below we briefly outline our methods; a more detailed description is available in online supplementary information.

To test how well the current guidelines safeguard long-term populations persistence, we used matrix population models to calculate 30-year projections of population sizes. We simulate seed harvesting as a reduction of the sexually produced new recruits. We generally modelled the most extreme scenario: the highest permitted seed fraction harvested every year. To allow comparison across species, we expressed effects of seed harvesting as relative population sizes, where *e.g.* 0.8 represents a 20% reduction of population size and 0.3 a 70% reduction over 30 years, in comparison to the population sizes that would be reached without seed harvesting (SM, section 4). As the effects of seed harvesting were independent of the biogeographic origins of the examined species (Table S2), we generally used all species in our dataset to test the guidelines of specific countries. We present the results separately for different growth types, as in the German the guidelines the recommended safe seed fractions are growth-type specific (16).

To find a better predictor of safe seed fraction than the growth types, we examined whether and which life history traits were better predictors of seed harvesting impacts (Figure 2). To enable practitioners to apply our findings, we restricted our analyses to five key life history traits readily available from public databases (21, 22, 36) or easy to estimate in the field: generation time, mean age at sexual maturity, the degree of iteroparity (frequency of reproduction) and clonality, and seed bank persistence (Figure 2, SM section 5). We then related these traits to the vulnerability of our 298 species to seed harvesting, defined as the slope of the relative decrease in population size with increasing seed harvesting (SM sections 3 and 6, Table S3).

To provide a quantitative basis for improving seed harvesting guidelines, we used generation time, the best predictor of species vulnerability to seed harvesting, to estimate safe seed fractions across species (SM section 7). The safe seed fractions were defined as the proportions of seed production where annual removal caused a <50% decrease of population sizes during 30 years of continuous seed harvesting, compared to the same populations without seed harvesting. A 50% decrease over 30 years corresponds to an annual decrease of about 2%. Importantly, this threshold ensures a >95% probability of population viability under environmental stochasticity in all analysed species but one (Figure S4).

To understand how environmental stochasticity affected our prediction for seed harvesting based on mean matrix population models, we simulated the effects of environmental stochasticity on population dynamics (SM section 8). This was possible in 108 species for which we had at least three spatial or temporal replicate matrix population models (so called individual models). We simulated environmental stochasticity as projecting population vector by randomly drawn individual matrix population models in each step, replicated 1000 times to obtain probability distributions of seed harvesting impacts. To understand how robust our estimates were to environmental stochasticity, we compared the safe seed fractions based on the mean matrix models to the respective medians of the safe seed fractions based on stochastic simulations (SM section 8.1). We also used stochastic simulations to test whether the thresholds of 50% population declines (see above) effectively prevented populations from extinction (SM section 8.2).

## Materials and Methods, Supplementary results

To quantify the effect of seed harvesting on wild plant populations, we used matrix population models (*35*). We first tested the impacts of seed harvesting by simulating the regulatory recommendations on seed harvesting in the wild of three regions where such regulations are in place (Australia, Germany and USA). Second, we calculated the population vulnerability to seed harvesting for each of the 280 plant species examined. Third, we related those effects to plant key life history traits (*i.e.* defining characteristics of their life cycles; e.g. generation time, age at maturity). In the fourth step, we used the life history traits that explained most of the vulnerability of natural populations to seed harvesting to formulate biologically-sound management recommendations. The ultimate goal of these recommendations is to introduce a threshold to seed harvesting so that (i) the population size does not decline more than by 50% over 30 years of consecutive (*i.e.* annual) seed harvest and (ii) the population may still have a 95% probability of persistence. All calculations and statistics were performed in R (*36*), and the reproducible, commented scripts are found as *Auxiliary material and will be available at Zonedo upon acceptance*.

### 1 Matrix population models

#### 1.1 General introduction

Matrix population models (MPMs, hereafter) are a widely used tool for investigating population dynamics (*35*). Briefly, an MPM describes the life cycle of an organism in terms of age, size and/or developmental stages along its life cycle and the transitions between stages, usually from one year to the next, as well as the sexual and clonal per-capita contributions to the population by individuals in each of those stages (Figure S1). One of the many applications of MPMs is to project the dynamics of a population through time (*35*), whereby a long-term population growth rate can be estimated (Figure S1). Importantly here, MPMs can also be used to calculate a wide range of population characteristics such as life history traits (*37*), extinction probability (*38*), and the effects of different hypothetical events (such as seed harvesting) on the long-term viability of a population (*39*, *40*).

In this study, we used MPMs to simulate seed harvesting as reduction of the per-capita contribution(s) describing seed production (Figure S1). We did so by simulating the harvesting of newly produced seeds while keeping all other demographic processes unaltered. The resulting MPM thus describes the population dynamics in a year where seed harvesting took place.

#### 1.2 COMPADRE database

We used data stored in THE COMPADRE Plant Matrix Database (version 5.0.0.), last accessed 25.8.2019 (*20*). In this version, COMPADRE contains 9121 MPMs from 647 published works describing life cycles of 760 plant species, ranging from algae to trees worldwide. MPMs in the database are accompanied by extensive metadata including the continent where the study was carried out, whether it was carried out in captivity or in the wild, and standardized information about each life cycle stage in three categories: propagules, individuals photosynthetically active, and individuals in vegetative dormancy. In the vast majority of MPMs in COMPADRE, the full MPM ***A*** is divided into three submatrices (*37*): ***U*** includes demographic processes that depend on survival of individuals alive at the beginning of the census (i.e., progressive growth, stasis, retrogressive growth, seed bank persistence, and vegetative dormancy), ***F*** includes sexual reproduction (e.g. production of seeds and juveniles), and ***C*** includes clonal reproduction (*i.e.* vegetative reproduction through ramets), such that

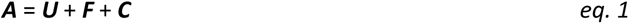

#### 1.3 Selection of the MPMs

We selected species and MPMs from COMPADRE based on the following criteria to allow for inter- specific comparisons to answer our questions:

- Only angiosperms and gymnosperms, since the ultimate goal of this study is to simulate the effect of seed harvesting on seed-producing plants.
- MPMs parameterised from field data from wild populations and under unmanipulated conditions, because the aim of this study is to understand the effect of seed harvest on natural, wild populations.
- MPMs for which the sexual reproduction component had been quantified explicitly, and separated from other processes in order to allow us to accurately perturb sexual reproduction (seed production; see below).
- MPMs that are irreducible, ergodic, and primitive, so the dominant eigenvalue (population growth rate) and other key properties could be calculated (*35*).
- When multiple studies per species were available (n = 235 species), we selected the single study per species that:
  - documented a seed bank, because inclusion of this transition in MPMs is vital to correct estimation of life history traits (*41*)
  - contained the highest number of individual MPMs (*i.e.*, from more populations or more years, see SM section 1.4) to use the most representative demographic information for the target species.

These selection criteria resulted in 467 MPMs from 467 plant species. Next, we checked the reliability of incorporating a seed bank in them or not. While survival of seeds in the seed bank is well documented in many demographic studies (*42*), between 42.9% and 47.3% of studies using MPMs in plant species unjustifiably exclude seed banks (8), thus assigning seedlings in year *t* to reproductive plants in *t*-1 (e.g. (*43*)). However, this assumption is only correct in species with a transient seed bank, i.e. seeds survive in the soil less than one year and thus, do not form a permanent soil seed bank (*41*). For those studies in our list where seed banks were not explicitly considered in their MPMs, we verified whether the species indeed have only a transient seed bank or not. We did so by carefully examining the original source of the MPM(s). If the source did not mention a seed bank, we further searched in the TRY database (*21*) for its potential existence. Consequently, we excluded 169 species where seed banks were unjustifiably excluded from their MPMs.

In twelve species, the simulated seed harvesting (SM section 2) did not cause any changes of population sizes, which suggests that generative reproduction was not correctly incorporated in these MPMs. We excluded these species from the further analysis.

This final selection criterion resulted in a dataset of 280 species (each with a representative MPM) from 83 plant families. This is the final set of species and data that were used for the simulations described below (**Error! Reference source not found.**).

#### 1.4 Mean MPMs vs individual MPMs

For the majority of studies in COMPADRE, MPMs are available for several annual transitions and populations. This was also the case in our final dataset. For all calculations, except in the case of stochastic simulations (section 8), we used a single *mean* MPM per species across all years and populations of demographic data available for that species. This mean MPM was calculated as the element-by-element arithmetic mean of the aforementioned MPMs, or pooled directly (e.g. weighted mean by sample size) from the individual-level data when provided by the author in the publication or through personal communications with the COMPADRE team.

For the stochastic simulations we used *individual* MPMs, which represented the population dynamics during a given annual transition and at a given population. We only used species that were represented in the database by at least three individual MPMs (section 8), resulting in 1578 individual MPMs from across 108 plant species in our dataset.

### 2 Simulating seed harvesting

We used the selected MPMs to simulate the impact of seed harvesting on populations. We first used the mean MPM (section 1.4) for each species, and simulated seed harvesting as a reduction in the values describing reproduction via seed in the sexual reproduction matrix ***F*** (see equation 1). Specifically, we created a modified MPM ***A’*** with reduced per-capita contributions of seed production in ***F***. To carry out our projections, we initiated the population vector ***n****_0_* as the stable stage distribution of the original MPM ***A***. This vector ***n****_0_* was obtained as the right-eigenvector of ***A*** following methods described by Caswell (2001). We then projected ***n****_0_* over 30 years using the modified MPM ***A’*** and the chain rule (*35*). We chose this period of time for our projections because it is long enough to observe even minor changes in the overall population size *N* that are not typically possible to quantify by short-term monitoring (*44*), while it is of sufficient length to fit within the active career of a land manager or conservation practitioner. We benchmarked the resulting population size *N*_30 harvest_ relative to the population size *N*_30 no harvest_ that would have been achieved in the absence of seed harvesting as in equation 2:

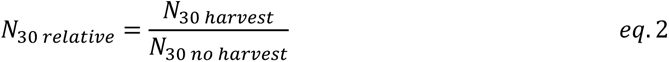

The relative population size *N*_30 relative_ thus ranges between 1 (when seed harvesting has no effect on population size; *N*_30 harvest_ = *N*_30 no-harvest_) to 0 (when the effect is so drastic it drives *N* to 0 within 30 years). For example, a value of *N*_30 relative_ = 0.1 means that the population size achieved with seed harvesting is 10% of the population size that would have been achieved without seed harvesting. The use of this metric as measure of seed harvesting impact allowed us to implement intra- and inter- specific comparisons, regardless of the variable population growth rates of each species’ population. When calculating the population sizes with and without harvest (*N*_30 harvest_ and *N*_30 no harvest_), we included only the active but not dormant (seed bank, dormant vegetative) life stages of the population vectors *N*_30_ because practitioners and scientists commonly evaluate population size based on counting active, standing individuals.

### 3 Vulnerability to seed harvesting

We used mean MPMs to calculate species vulnerabilities to seed harvesting. For each species, we created 101 MPMs that describe the population dynamics when harvesting 0-100% of seed production, in 1% steps (Figure S1). As in section 2, we used the virtual MPMs to project population sizes over 30 years. We then fitted an exponential-decay model to quantify the effects of the varying proportion of harvested seed (p) on the relative population size in 30 years (*N*_30 relative_) as follows:

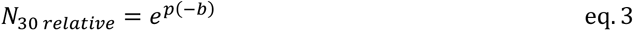

where *b* determines how steeply the relative population size (*N*_30 relative_) decreases with increasing proportion of harvest pressure, such that the larger *b*, the steeper this decrease is. We refer to this coefficient as *vulnerability to seed harvesting* (**Error! Reference source not found.**).

### 4 Testing current recommendations

Next, we used MPMs to simulate the impact of seed harvesting according to the current rules on the relative population size *N_30 relative_*. As far as we are aware of, explicit recommendations for the maximal proportion of seeds that can be harvested from natural populations so far exist only in three countries. In USA and Australia, this value is 20% and 10%, respectively, for common plant species when harvesting seeds for restoration projects (*13*, *14*). German rules are available for herbaceous plants: 2% for annual and 10% for perennial species when harvested every year (*15*).

As the current recommendations are partly growth-form specific (*15*), we examined the reduction in relative population size as a function of plant growth form: annuals, herbaceous perennials, epiphytes, lianas, palms, succulents, shrubs, and trees, as indicated in the COMPADRE metadata. We excluded growth forms represented by less than 5 species: epiphytes (n=4) and lianas (n=1), as well as plant species whose generation time disagreed with the metadata of the species, in particular annual species with generation times larger than two years (n=4). As the vulnerability to seed harvesting of individual species (Section 3) depended neither on a continent nor on the interaction between a continent and plant growth form (Table S2), we grouped species only by growth form and used the same set of species to test the recommendations from Australia, USA and Germany (Figure 1 in the main text).

### 5 Life history traits

We used life history traits to explain species vulnerability to seed harvesting. A life history trait is a key feature that describes the life cycle of the organism (e.g. generation time, survival of seeds in the seed bank, clonal propagation). As our ultimate motivation was to facilitate the translation of our findings to land managers and practitioners, out of the wide range of life history traits that can be derived from MPMs (e.g (*35*, *37*)), we selected the traits that are readily available in trait databases or easy to estimate in the field (Table S3). All life history traits were calculated based on the matrix A of the mean MPM of each of our 280 species.

### 6 The effect of life history traits on vulnerability to seed harvesting

We used linear models to determine which life history traits (generation time, degree of iteroparity, age at sexual maturity, seed bank residence, clonality) best explained species’ vulnerability to seed harvesting (Section 3). We also added plant growth form as an explanatory variable (as defined in the COMPADRE database (*20*)) to the model to test whether it explains any additional variability. Restricting the model to key life history traits allowed us to keep the full model and avoid model selection, which is known to produce exaggerated effect sizes and spurious effects (*45*). Species vulnerability to seed harvesting was log-transformed prior analysis to achieve normality. Other explanatory variables except plant growth type (factor) were log-transformed and standardised to adhere to the model assumptions of normally distributed errors.

To illustrate the importance of the life history traits for predicting the species vulnerability to seed harvesting (Figure 2 in the main text), we expressed the relative importance of each predictor in the model as the proportion of explained variability assigned to each predictor. As the explained variability can depend on the sequential order of the predictors in the model, we averaged the explained variability for each predictor across all possible ordering of the predictors using the R package *relaimp* (*46*). To visualize effect sizes of the effects of life history traits on species vulnerability to seed harvesting, as well as uncertainty of these effects, we used 95% credible intervals, a Bayesian analogue of confidence intervals. These were calculated from 10,000 simulations of the mean and variance of each estimate, using the *sim* function in the R package *arm* with non-informative prior (*47*).

We also ran a model including the phylogenetic relationships among species to test the extent to which the explanatory power of life history traits on species’ vulnerability to seed harvesting is in fact driven by the phylogenetic inertia in plant life history traits (*48*). We used a phylogenetic generalized least square model to include the phylogeny of our species. We obtained the phylogeny from COMPADRE, following methods detailed elsewhere (*37*). With this model, we estimated Pagel’s *λ* (not to be confused with the population growth rate, also referred to as *λ* in the demographic literature (*35*)), a measure of phylogenetic signal in the trait structure. Briefly, Pagel’s *λ*=0 indicates no effect of the phylogenetic structure in the dataset in explaining variation in a given trait, while Pagel’s *λ*=1 indicates that the phylogenetic structure perfectly predicts, i.e. is responsible for, the life history trait structure.

Negative values suggest that closely related species have more different traits than would be expected by chance ((*48*). We found that the phylogenetic signal was overall weak and negative (Pagel’s *λ*=-0.1). Based on this result, we opted to present in this paper results from the linear model without phylogenetic correction.

### 7 Assessing limits of seed collection

We used the mean MPM per species to estimate what fraction of seed production one can collect from a natural population while only moderately affecting its dynamics. As a moderate effect we defined a reduction in population size *N* to not below 50% of the size that would have been achieved without seed harvesting during 30 years of a constant annual harvest intensity. While a reduction of population size by up to 50% over 30 years may seem relatively high, it corresponds to an annual decline of <2%. This threshold also allows for the persistence of the natural population under environmental stochasticity in >99% of species (see section 8.2).

For each species’ MPM, we simulated the effect of seed harvesting as a reduction of seed production transition by 0-100%, in 1% intervals. We used such reduced, virtual MPMs to simulate population dynamics across 30 years, and we recorded the final population size and expressed it as relative to population size that would be achieved without seed harvesting (see section 2, note this calculation is the same as the first step of the calculation of vulnerability to seed harvesting, Section 3). Besides annual harvests, we also modelled the effect of harvesting seeds every 2, 5 or 10 years because reducing harvesting frequency up to once in 10 years is sometimes recommended to limit negative effects of seed harvesting on population dynamics (*11*). In this case, we modelled population dynamics with the original mean MPM while the reduced MPM was used every 2^nd^, 5^th^ or 10^th^ run. As the safe fraction for seed harvesting, we considered the largest proportion of seed that was possible to harvest without exceeding the 50 % reduction of the relative population size.

We related the safe fraction for seed harvesting to the generation time of plants – the most important predictor of species vulnerability to seed harvesting, which alone explained 52.3% of total variability in species vulnerability to seed harvesting. We used non-linear regression in *R* (*nsl*) to describe the sigmoid relationship between the safe fractions of seed harvesting and the generation time, and used function *PredFit* in package *investr* (*49*) to generate confidence intervals for the relationship (Figure 3 in the main text).

### 8 Effect of environmental stochasticity

In a subset of our studied species, we simulated the effects of environmental stochasticity on population dynamics to understand how environmental stochasticity affects our prediction for seed harvesting based on mean MPMs. We used all species in our dataset represented by at least three individual MPMs (section 1.4), resulting in 1,578 MPMs across 108 plant species. We simulated environmental stochasticity as projecting vector of stable stage distribution of the mean MPM by randomly drawn individual MPM in each step. To obtain a probability distribution of results under environmental stochasticity, we repeated this process 1,000 times. We expressed the results as *N*_30 relative_ (equation eq. 2). The effects of seed harvesting were simulated as above (section 4), with the difference that in each of the 30 annual time-steps in each of the 1,000 simulation runs, we randomly drew an individual MPM from the set of individual MPMs available for a given species.

#### 8.1 The effect of seed harvesting on population size based on environmental stochasticity versus mean MPM

To understand how environmental stochasticity affected our results, we estimated the robustness of our results in stochastic environments. As an example, we used the effect of harvest of 20% of seed production, expressed as *N*_30 relative_, and simulated seed harvesting either using mean MPMs or stochastic simulation. We then compared the safe fraction for seed harvesting (*N*_30 relative_ > 0.5) based on the mean MPMs to the median of safe seed fraction based on the stochastic simulations.

The median of relative population sizes *N*_30 relative_ based on 1,000 permutations of stochastic simulations (y axis in Figure S3) closely correlated with the *N*_30 relative_ based on the mean MPMs. Interestingly, the relative population size *N*_30 relative_ based on stochastic simulation (orange points in Figure S3) was slightly higher than the *N*_30 relative_ based on mean MPMs (black line in Figure S3), especially in species that are more vulnerable to seed harvesting. Consequently, the safe fraction for seed harvesting based on the median of stochastic simulations was on average 0.017 higher that safe fraction based on the mean MPMs (Figure S4). This suggests that environmental stochasticity partly buffers the predicted decrease of population size caused by seed harvesting.

#### 8.2 Threshold for seed harvesting based on mean MPM versus extinction probability

In the models above, we set a threshold for seed harvesting so that the relative population size *N_30 relative_* decreased not below 50% of the population size that would be achieved without seed harvesting. In this section, we tested whether this threshold also prevented populations from extinctions. For each species, we computed what fraction of seeds could be sustainably harvested without causing extinction in at least 95% of stochastic simulations. We considered a population to go locally extinct when *N_30 relative_* < 0.01 (see section 2 for definition of *N_30 relative_*). For each species, we compared the threshold based on the 95% probability of population survival with the threshold based on mean MPM and *N_30 relative_* > 0.5.

In the vast majority (>99%) of examined species, the threshold based on *N_30 relative_* > 0.5 (as calculated using mean MPMs, black line in the Figure S5) allowed for the collection of a lower proportion of seeds than the threshold based on 95% probability population survival when using stochastic simulations (individual points in Figure 5). This suggests that the rules based on *N_30 relative_* > 0.5 derived from the mean MPMs prevent populations from going locally extinct.

---

**Figure S1:**
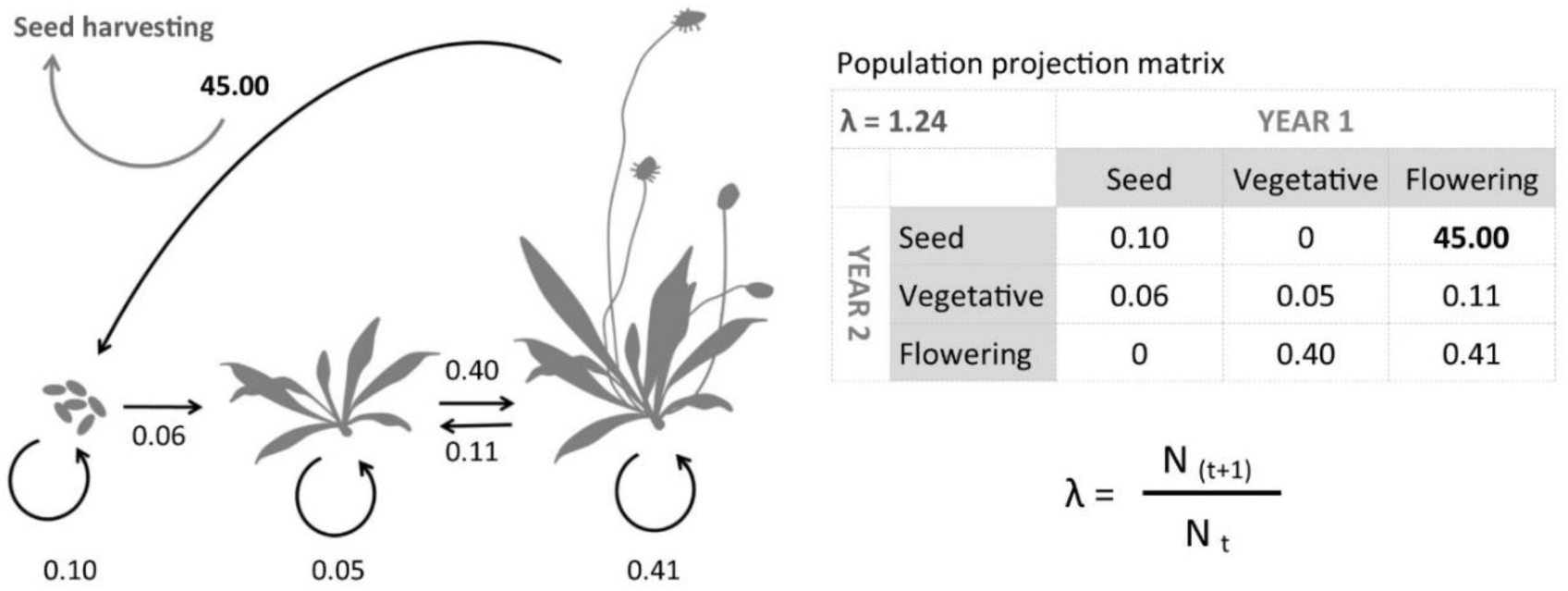
Life cycle of a hypothetical plant species with three stages (seedbank, juvenile, and adult) and its corresponding matrix population model (MPM), with *λ* indicating its long-term population growth rate, which is a function of population size (*N*) between two time-points *t* and *t*+1. Seed harvesting in this study was simulated by manipulating the transitions that describe generative reproduction.

**Figure S2.**
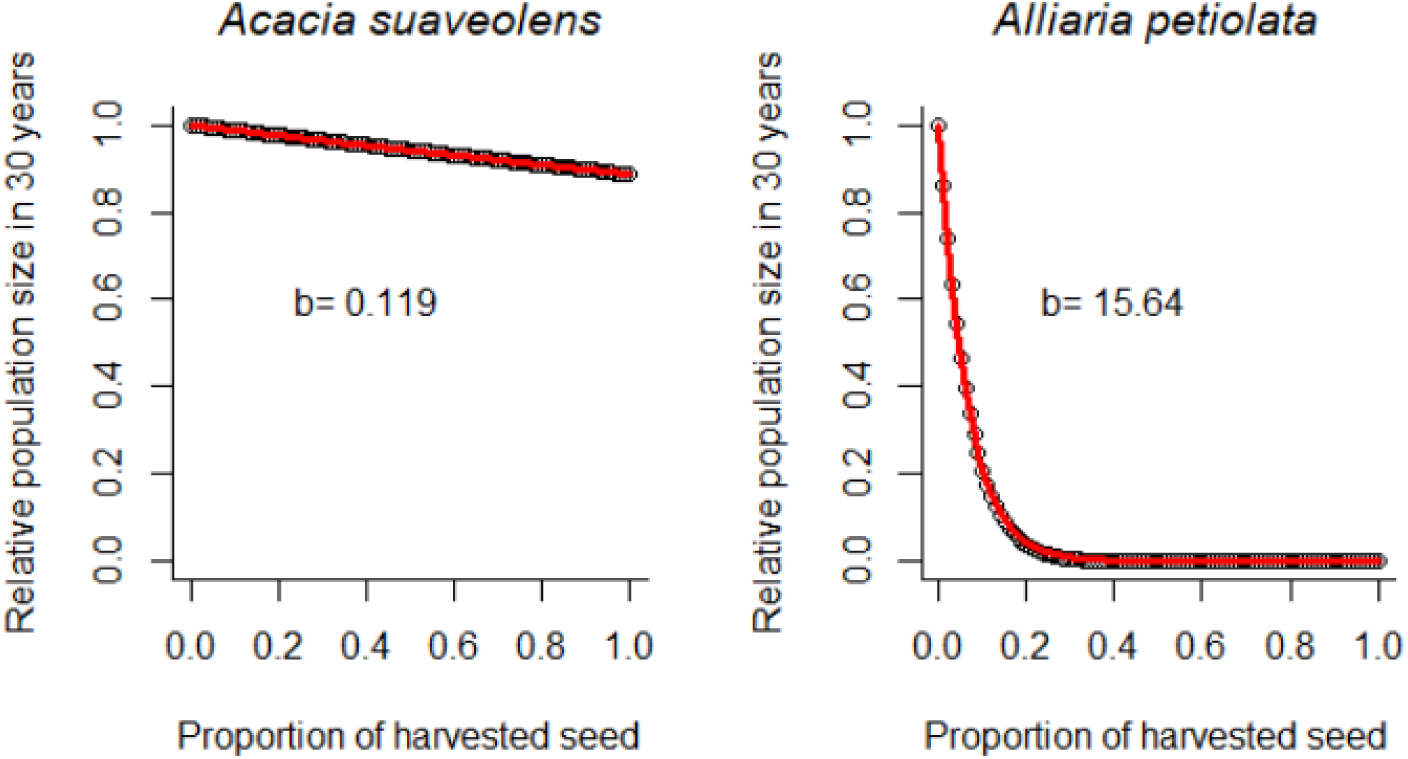
Vulnerability of population dynamics to seed harvesting (*b* in equation S3) in two of our 280 examined plant species. Note how the larger the value of *b*, the more vulnerable the given species is to seed harvesting. Black dots: simulated values; red line: fitted exponential-decay model as per equation 3.

**Figure S3:**
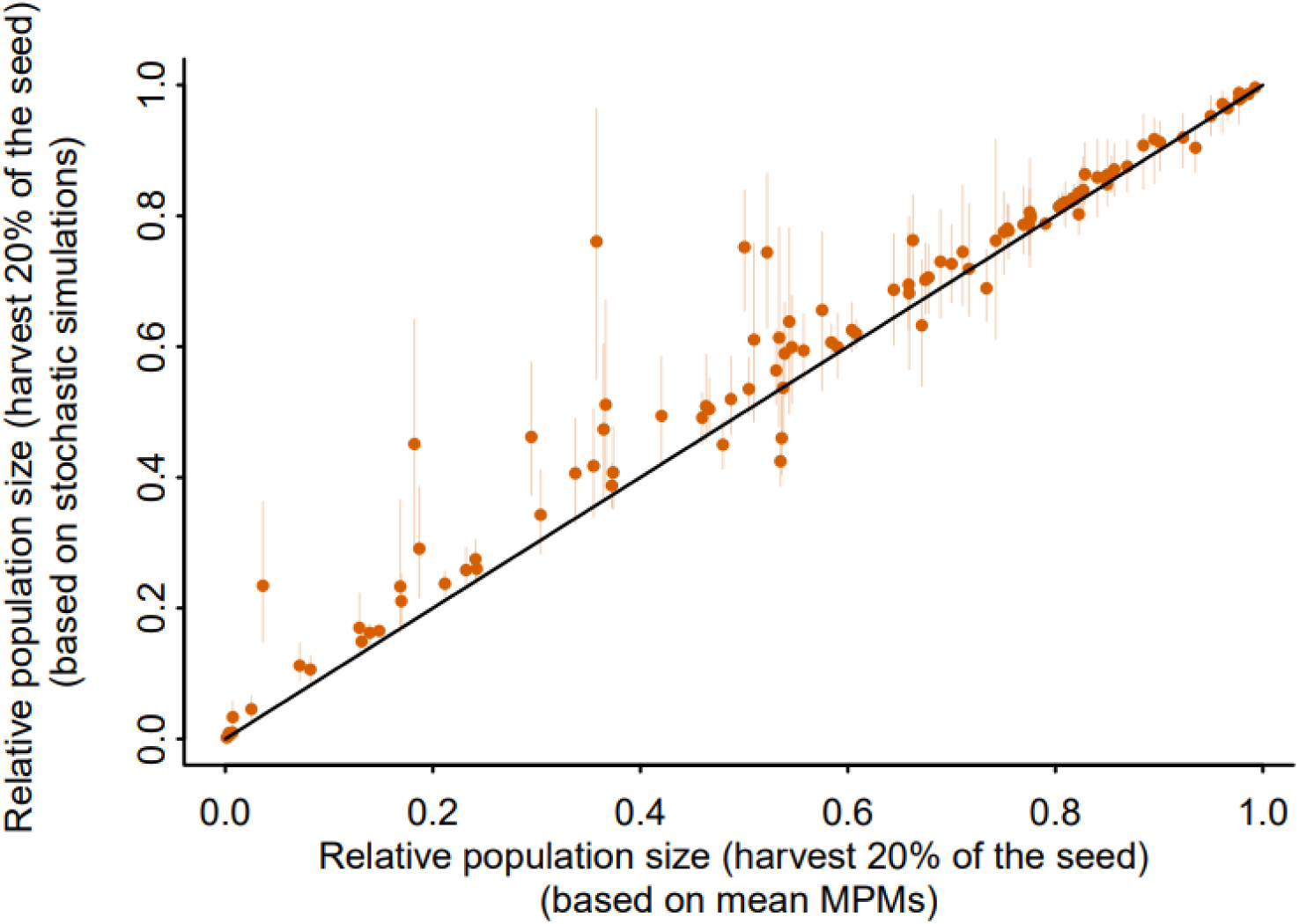
Comparison of relative population sizes (*N_30 relative_*) when 20% seeds were harvested based on mean MPMs (x axis and the 1:1 black line) versus from calculations with environmental stochasticity (y axis).

**Figure S4:**
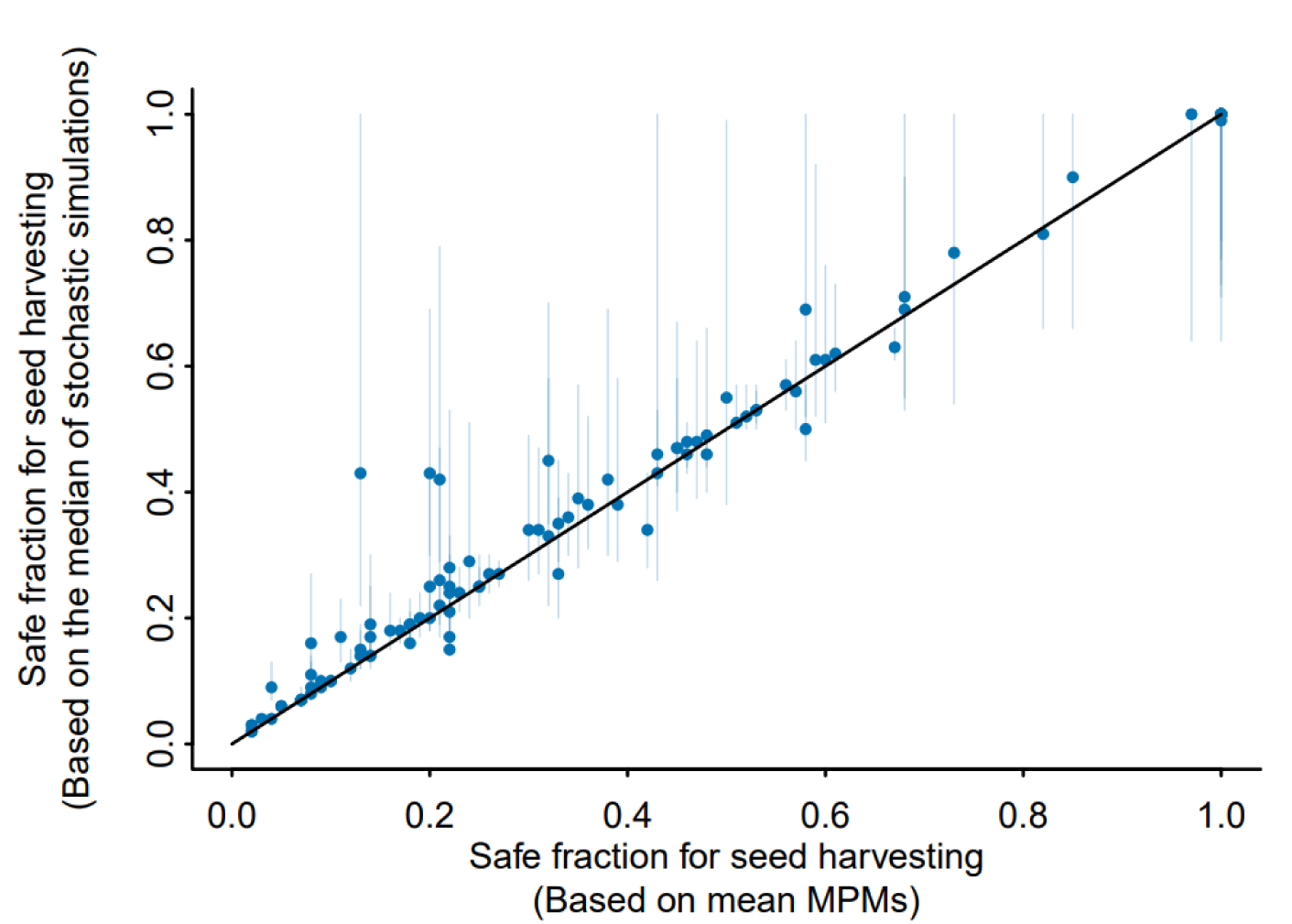
The safe fractions for seed harvesting based on *N_30 relative_* > 0.5 as calculated from the mean MPM (x-axis and 1:1 black line) versus the same safe fraction based on stochastic simulations (with 95% CI).

**Figure S5:**
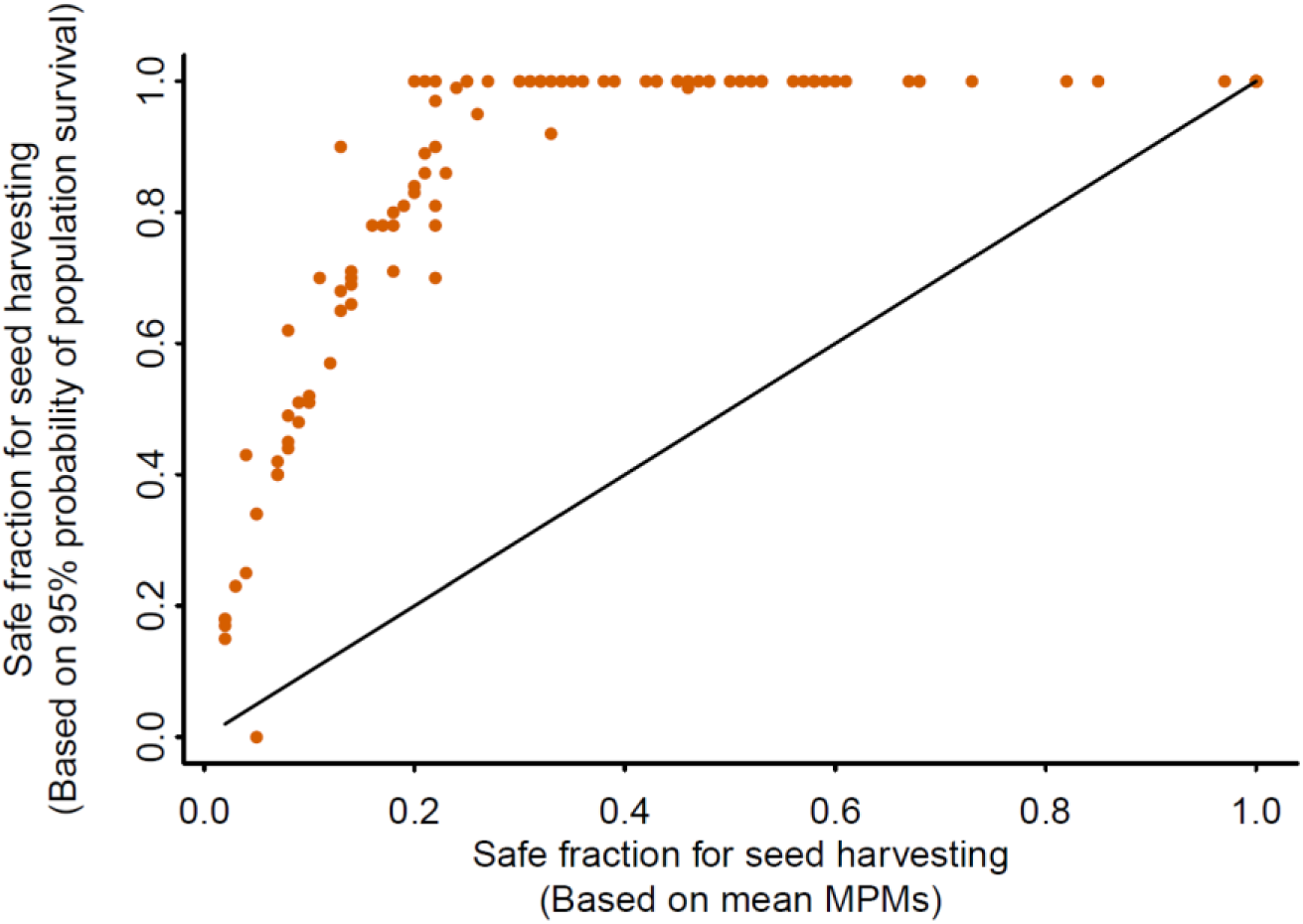
Comparison of the threshold for maximal seed harvest based on *N_30 relative_* > 0.5 as calculated from the mean MPM (x-axis and the 1:1 black line), with the maximal seed harvest that allows 95% probability of population survival of each considered species, as based on stochastic simulation.

**Table S1:**
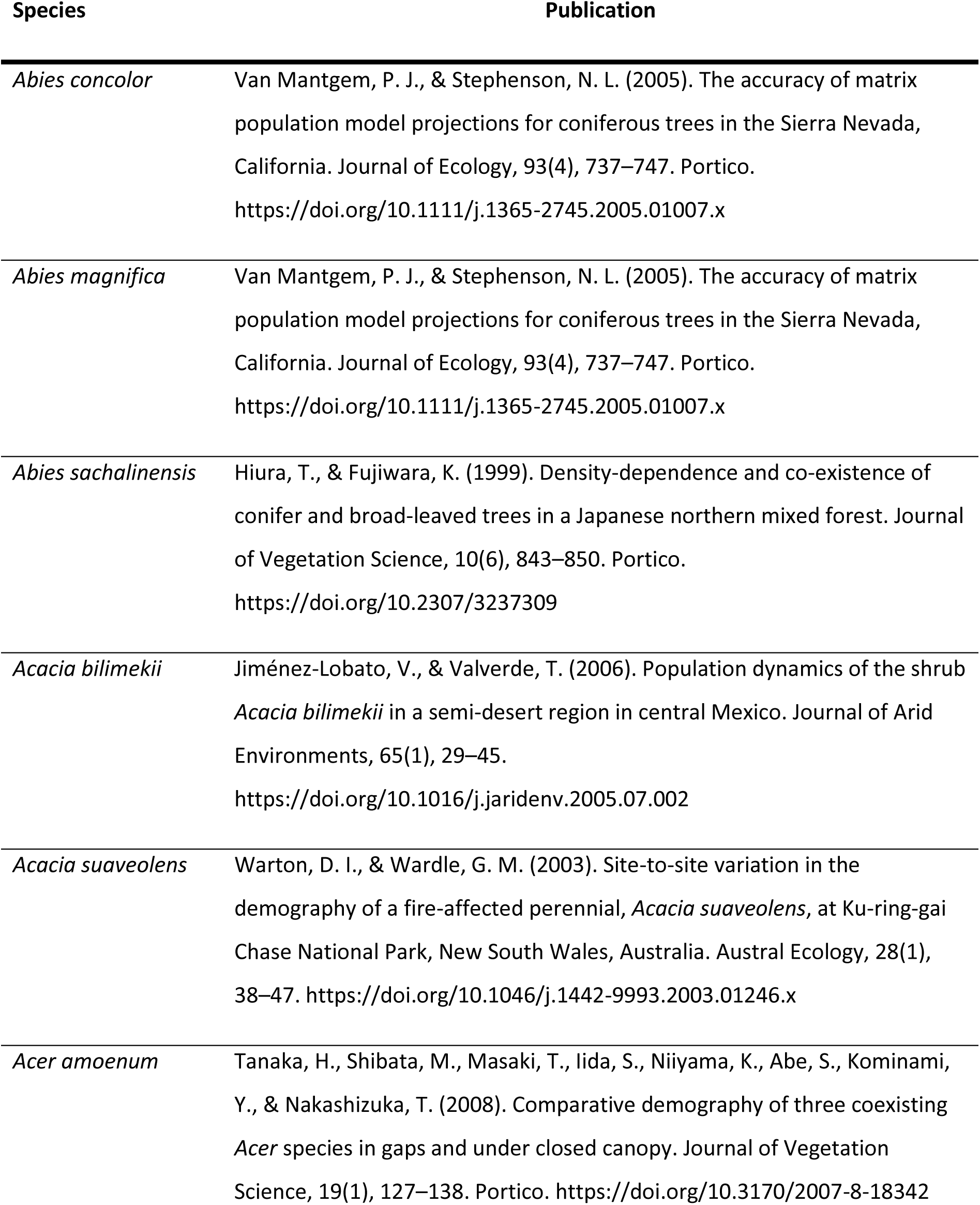
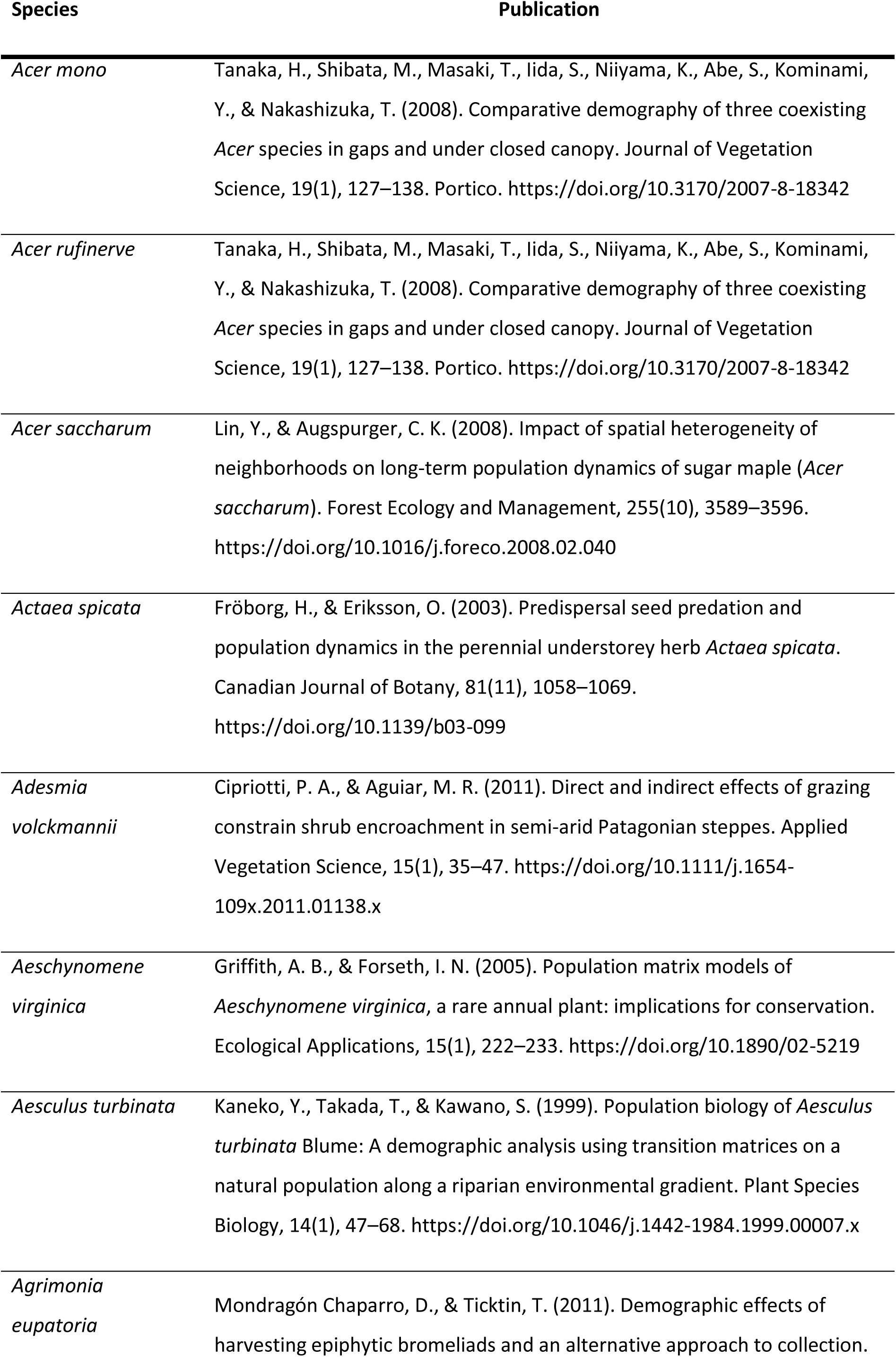
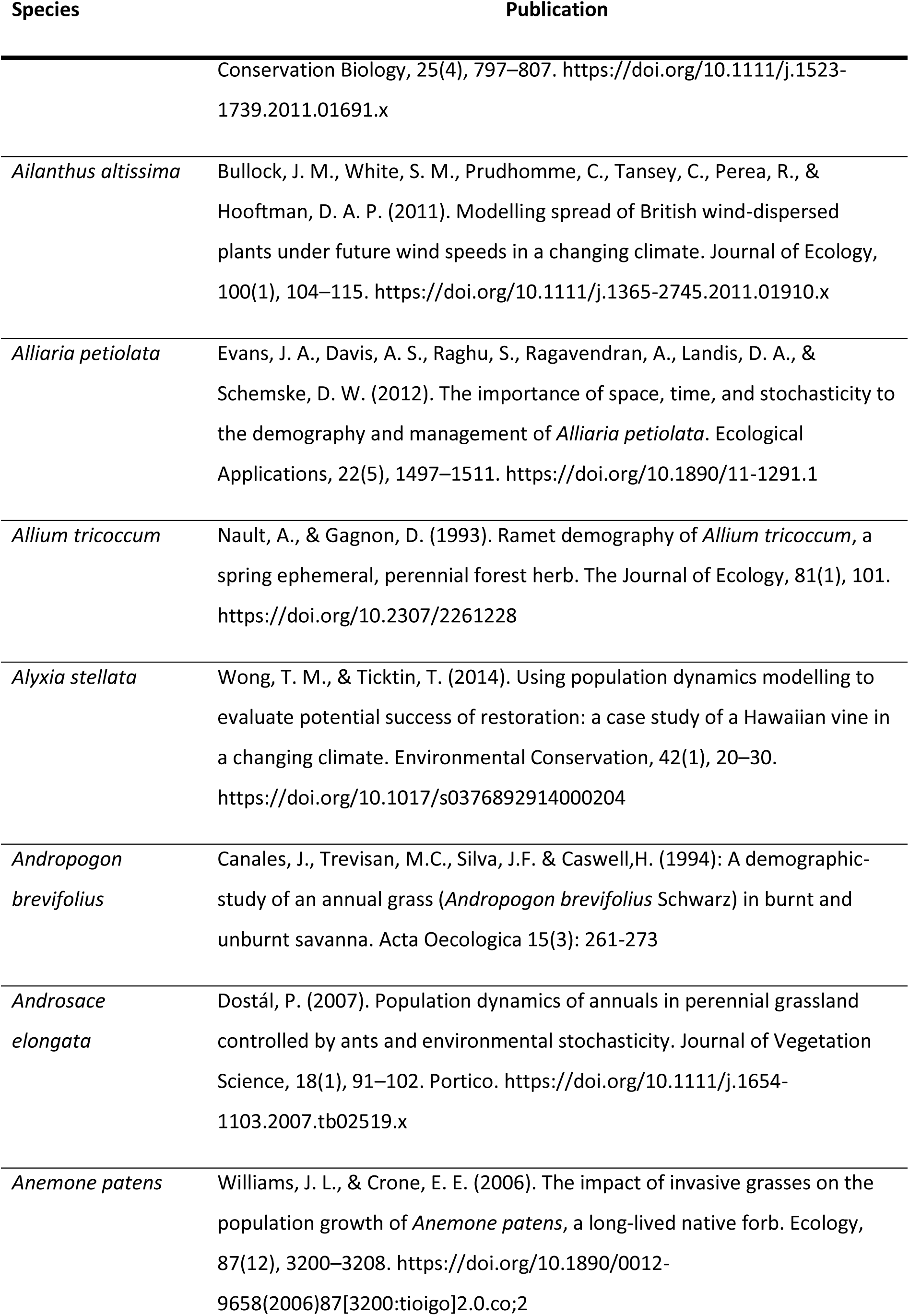
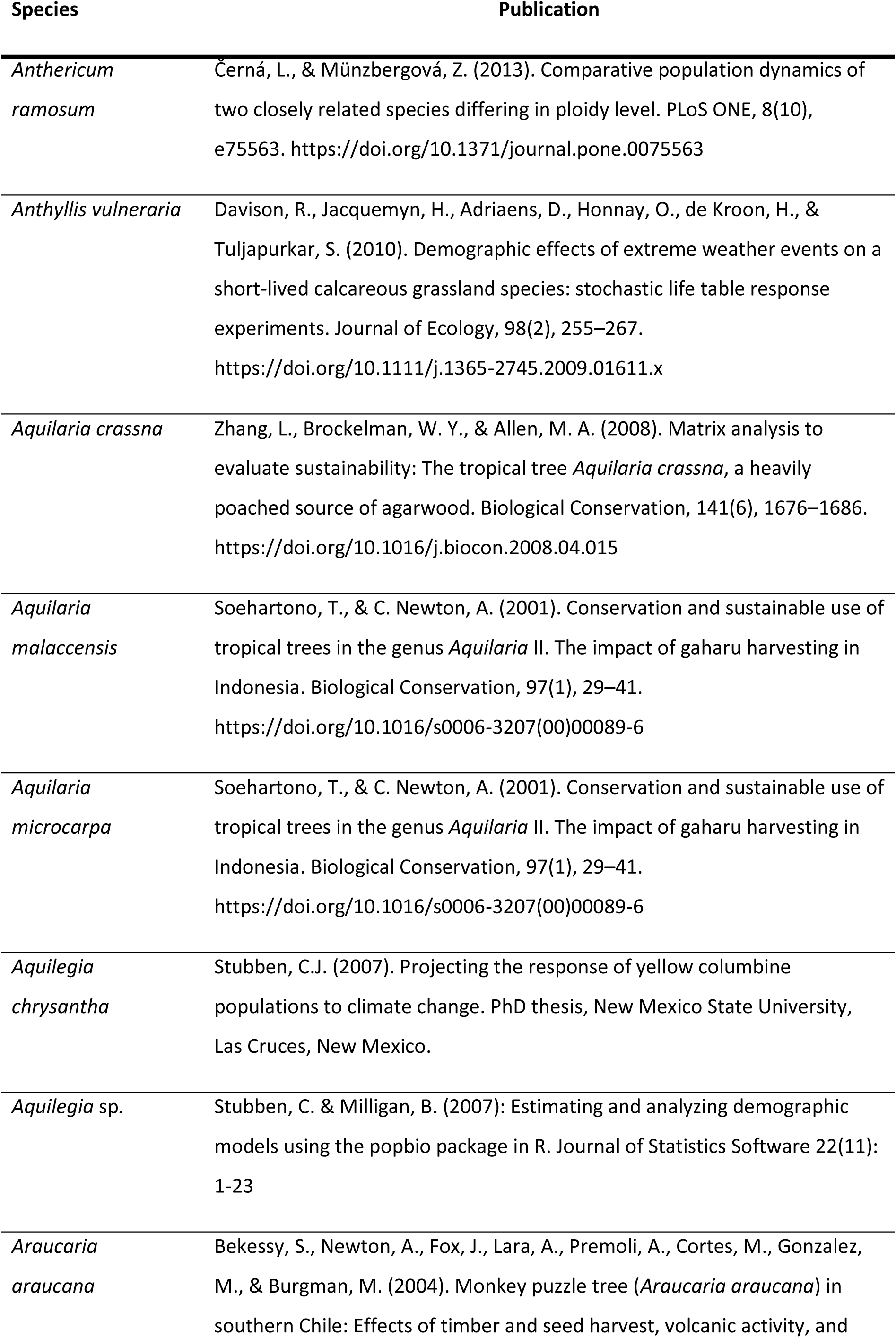
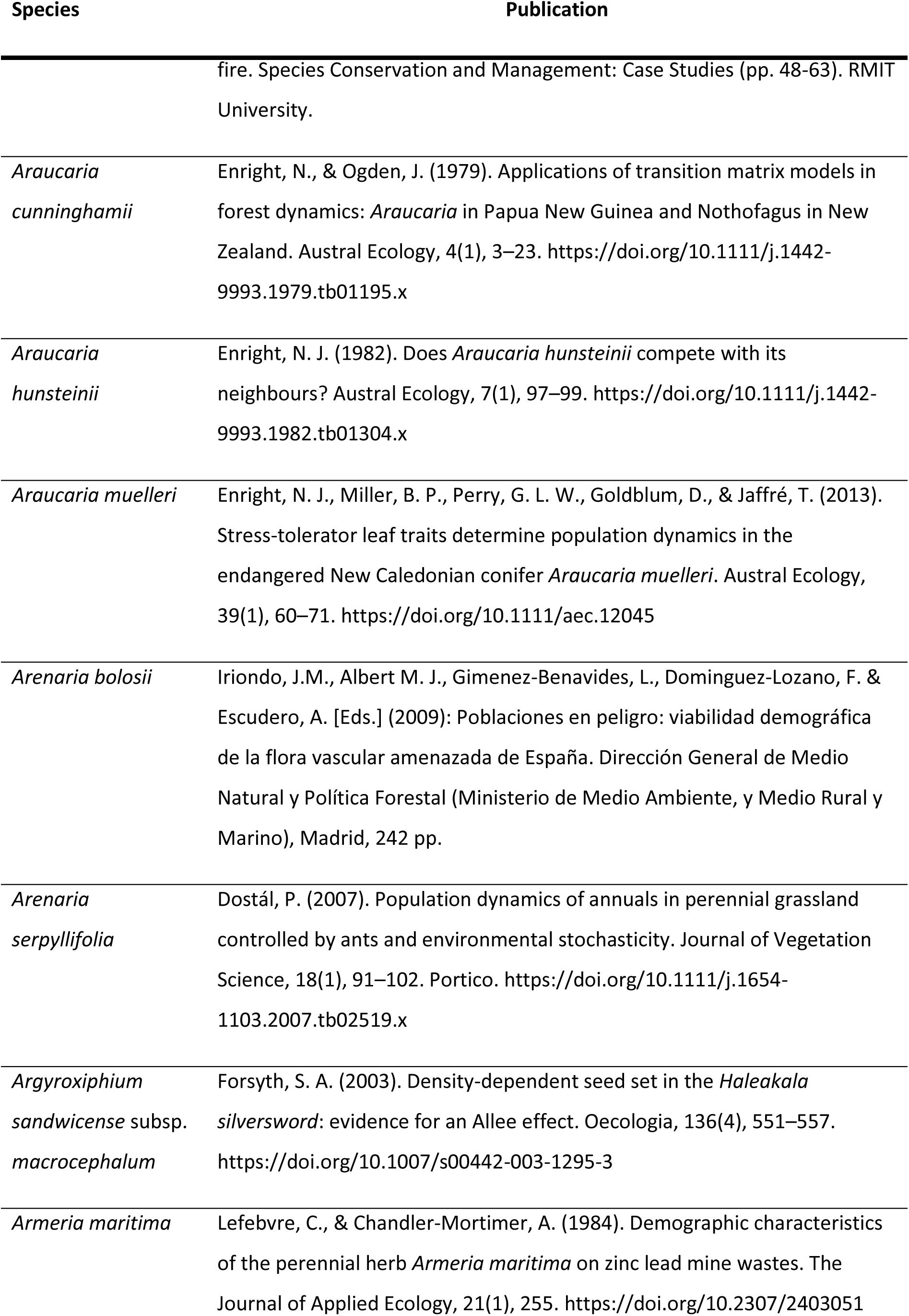
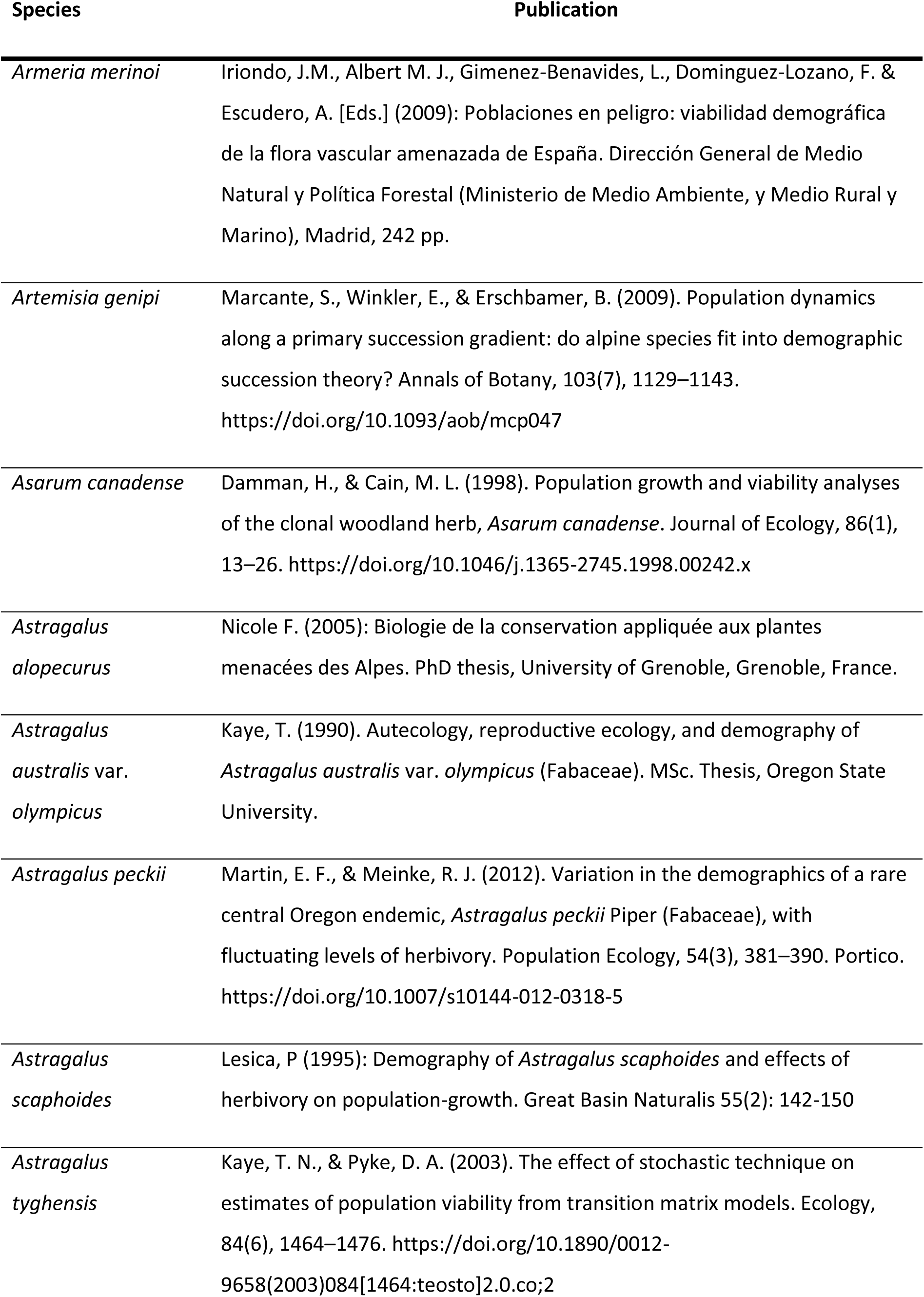
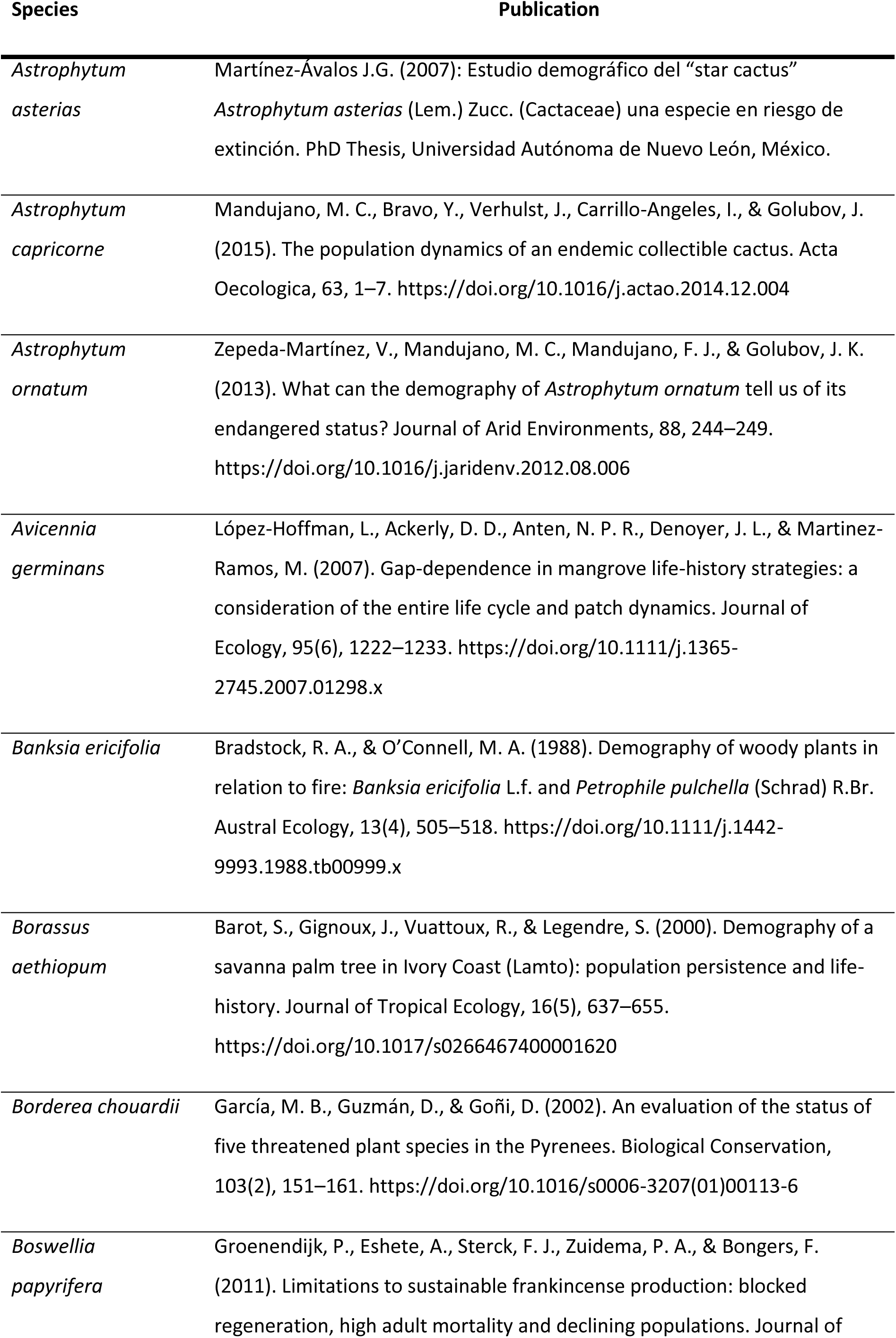
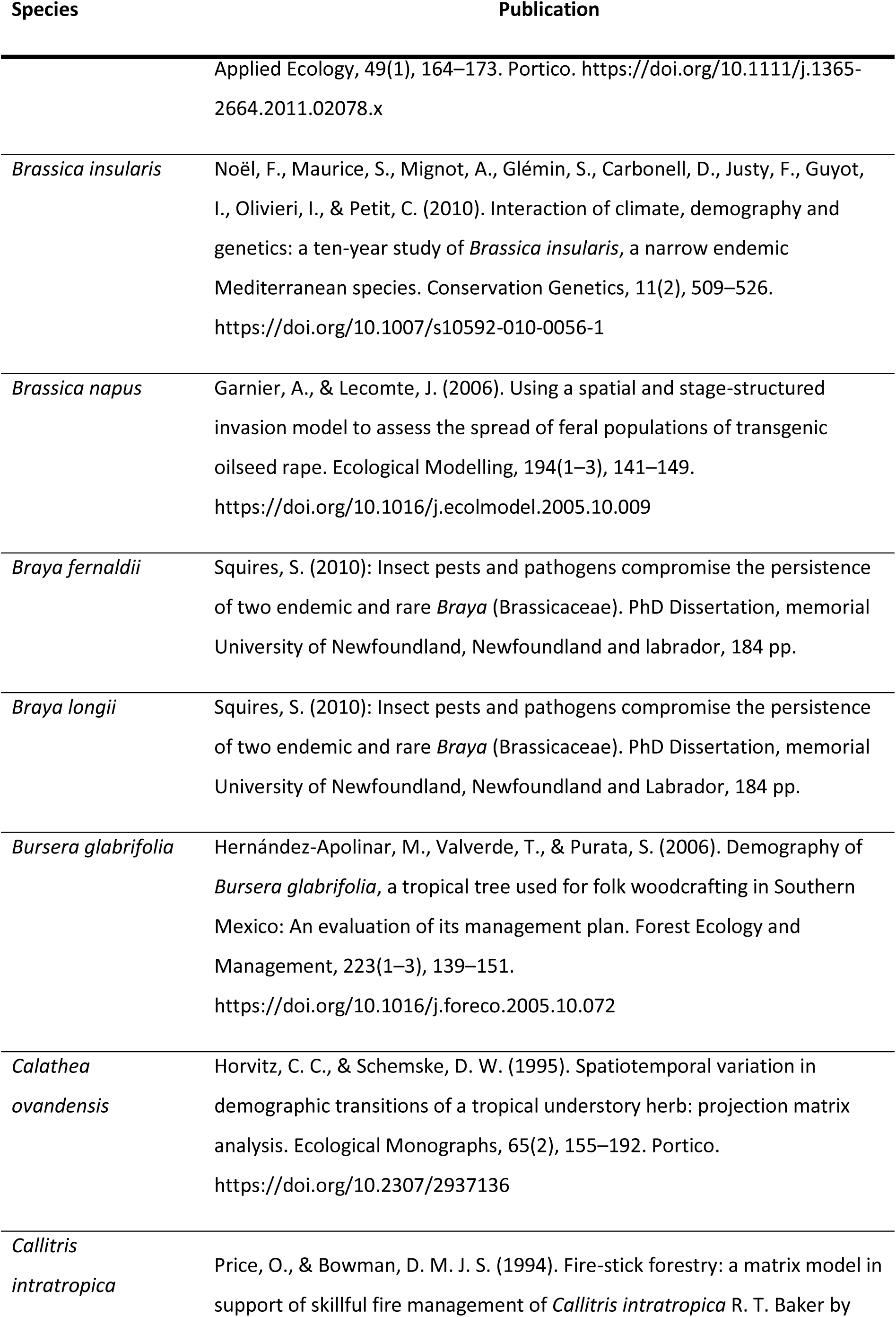
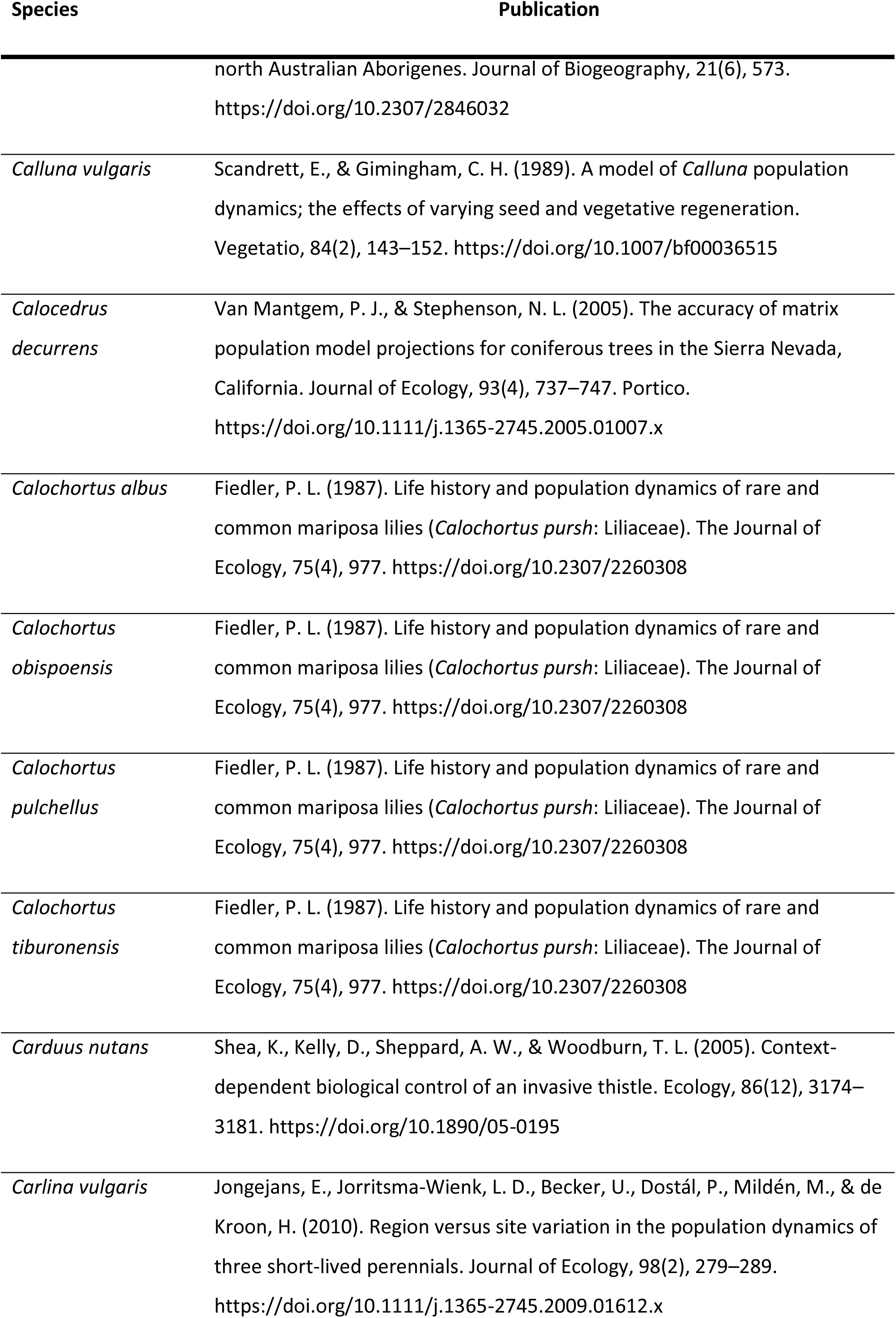
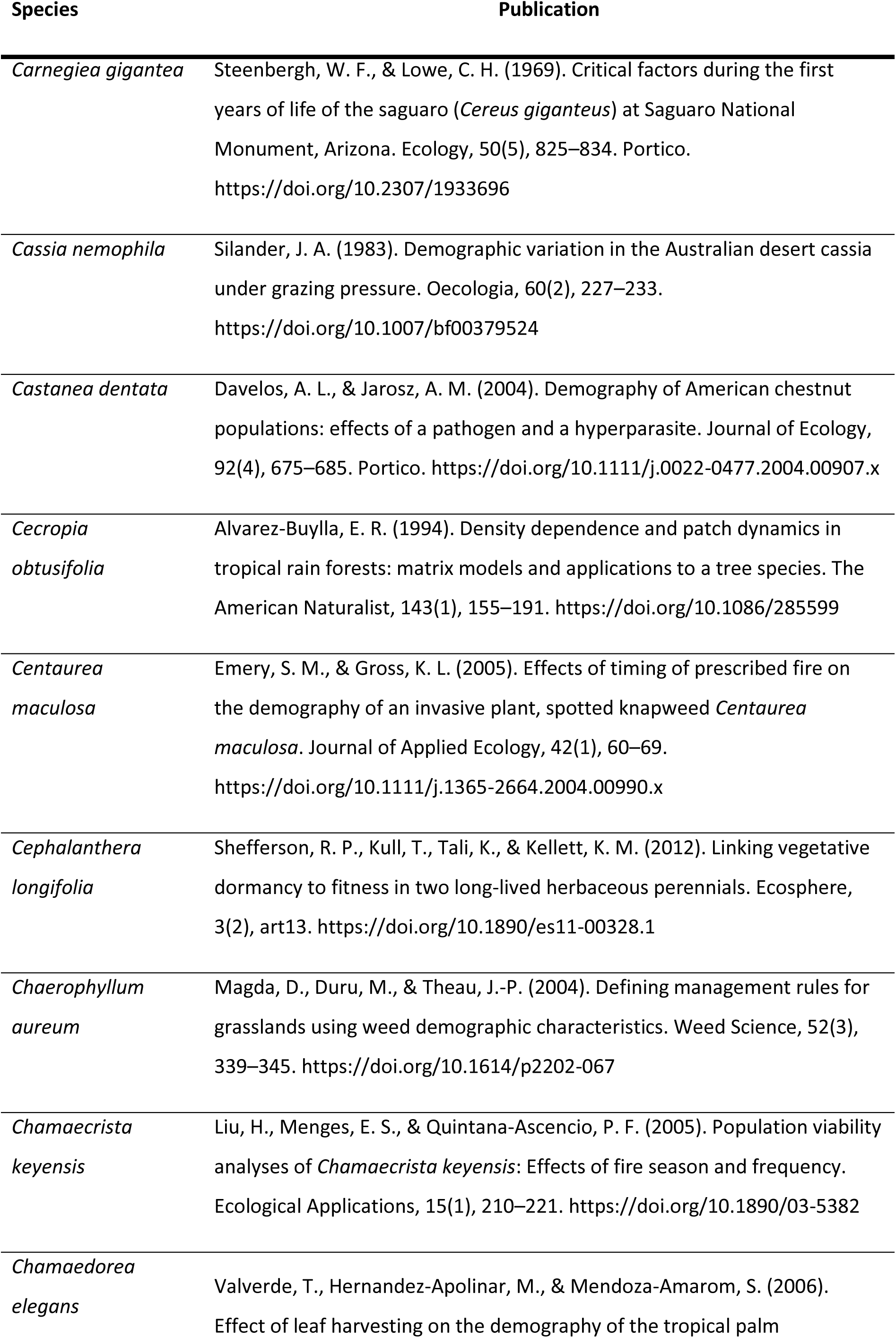
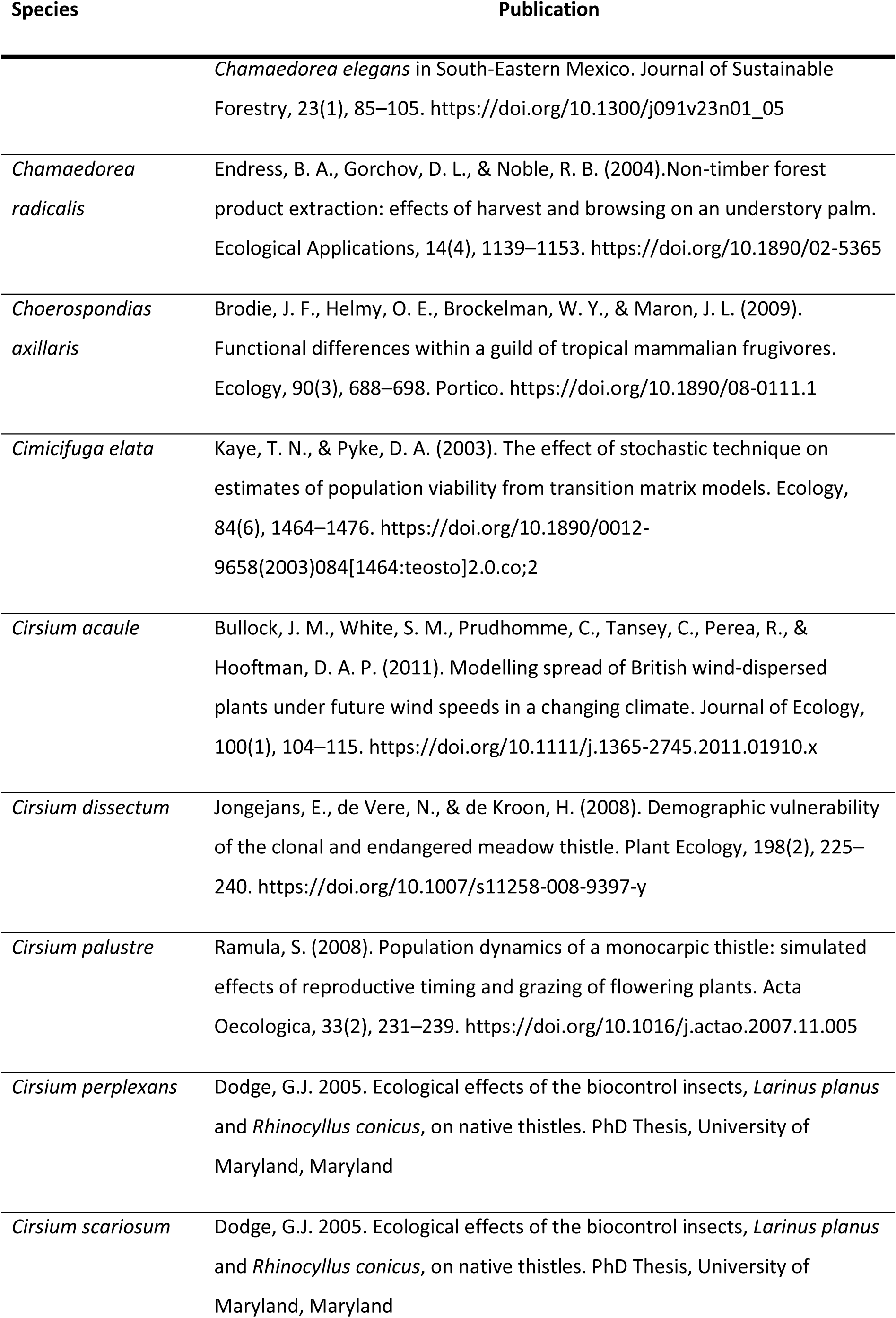
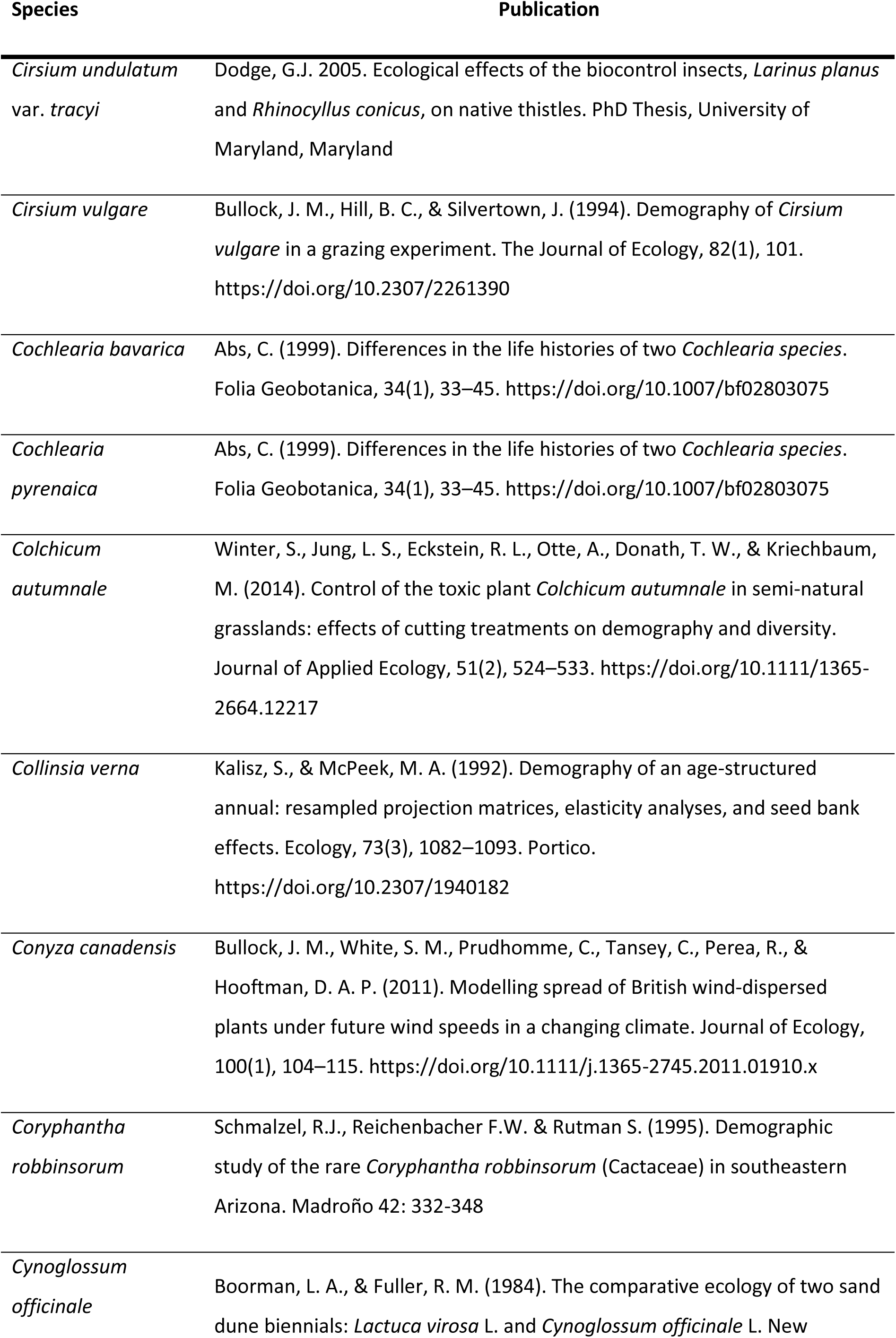
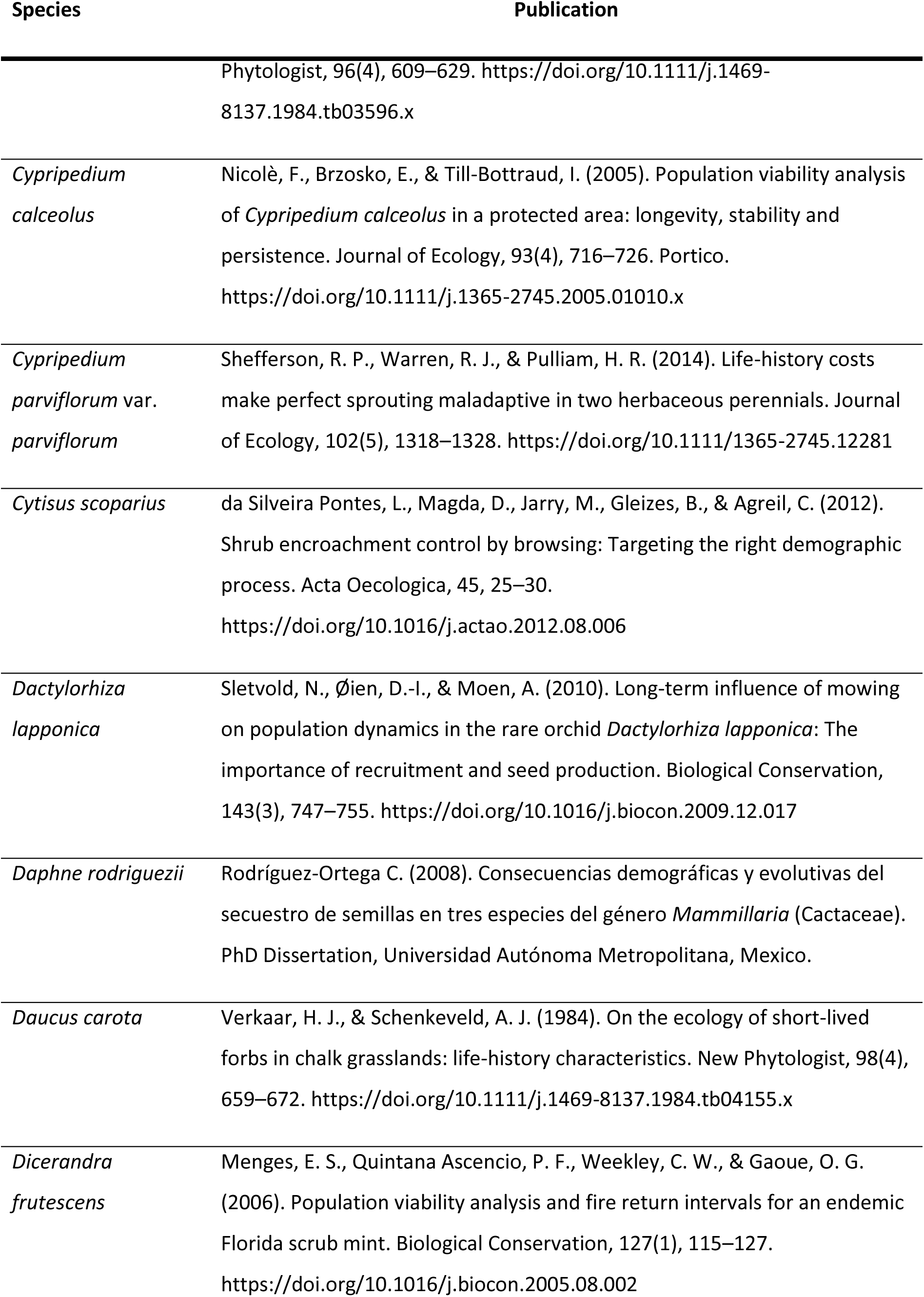
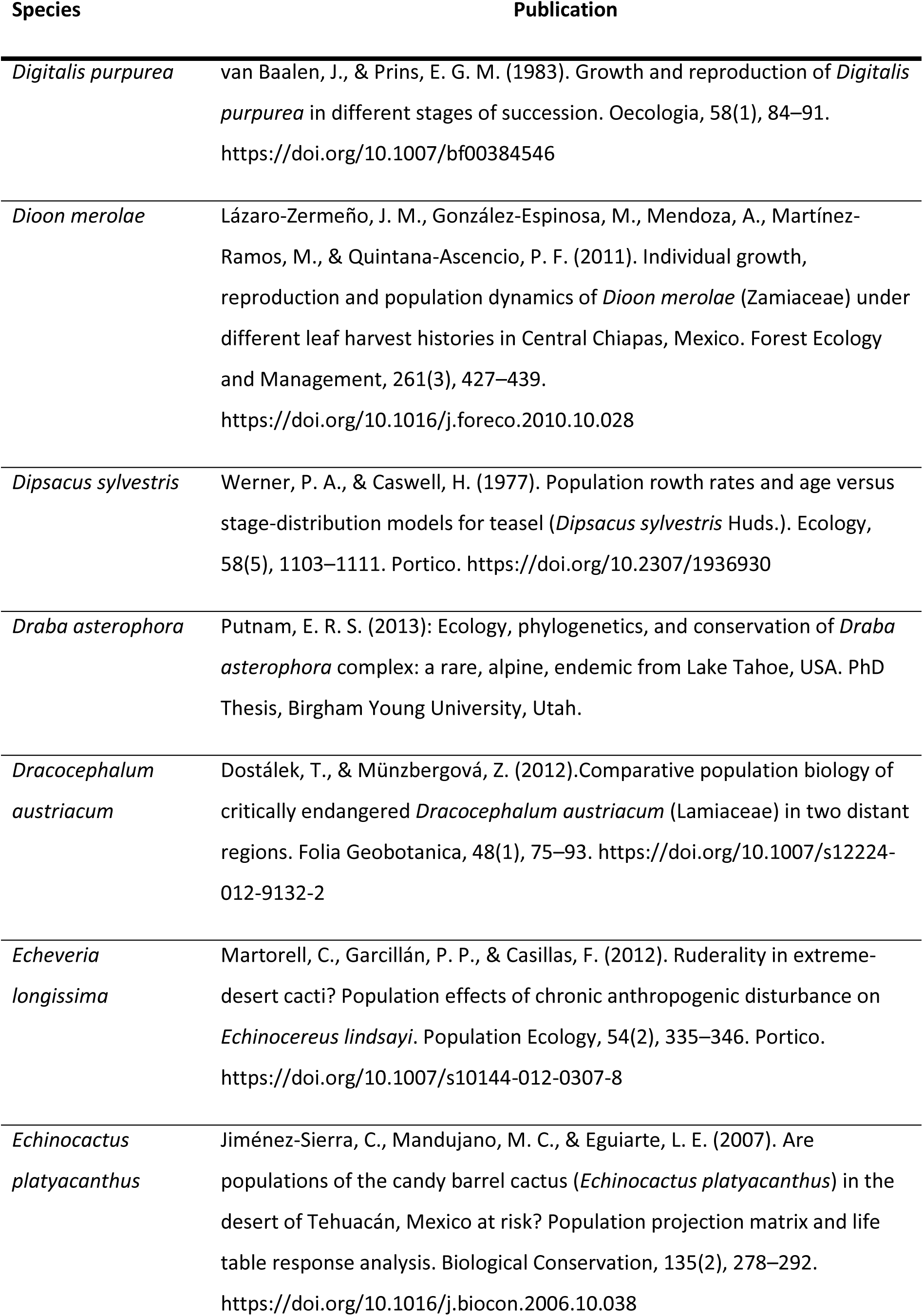
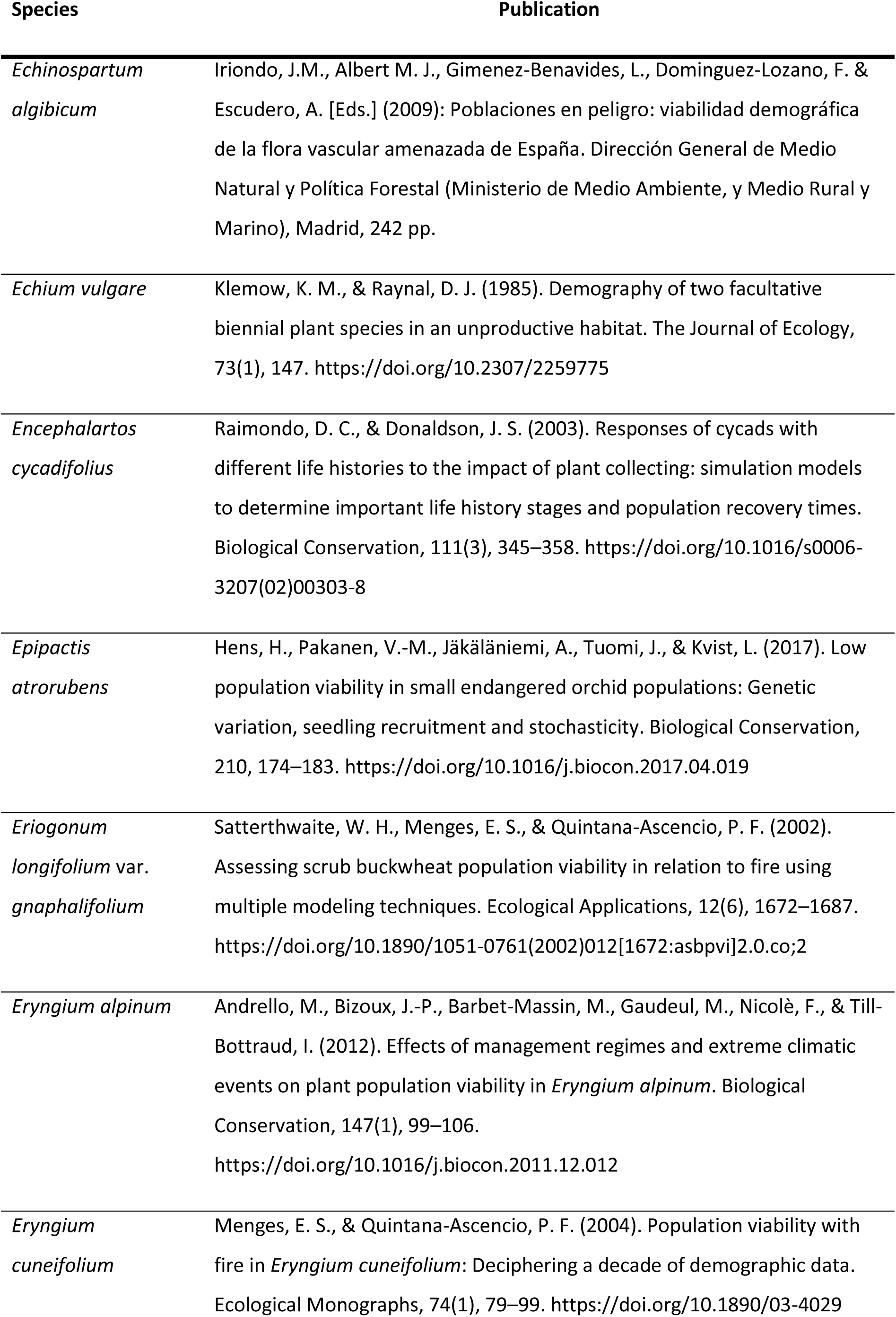
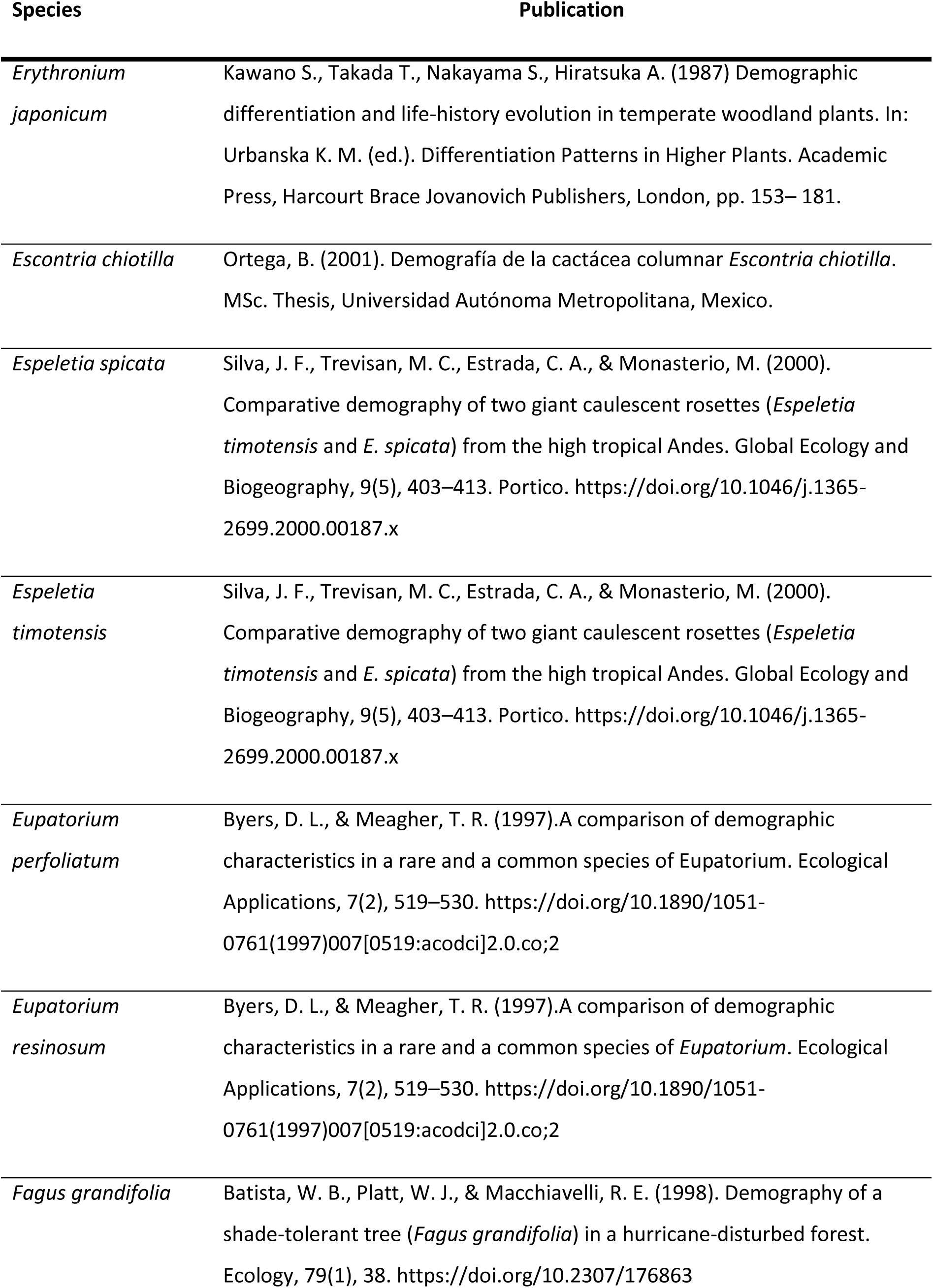
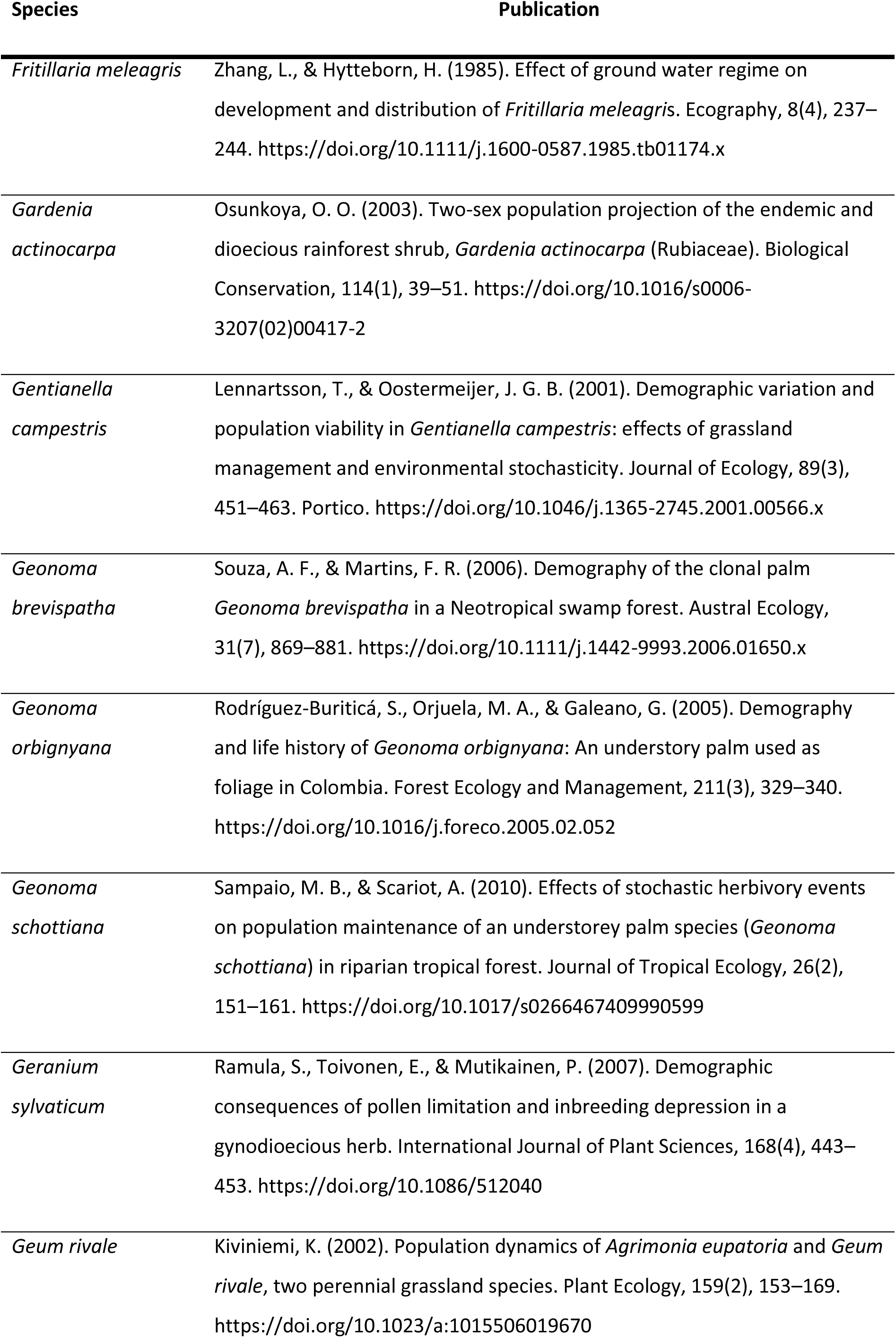
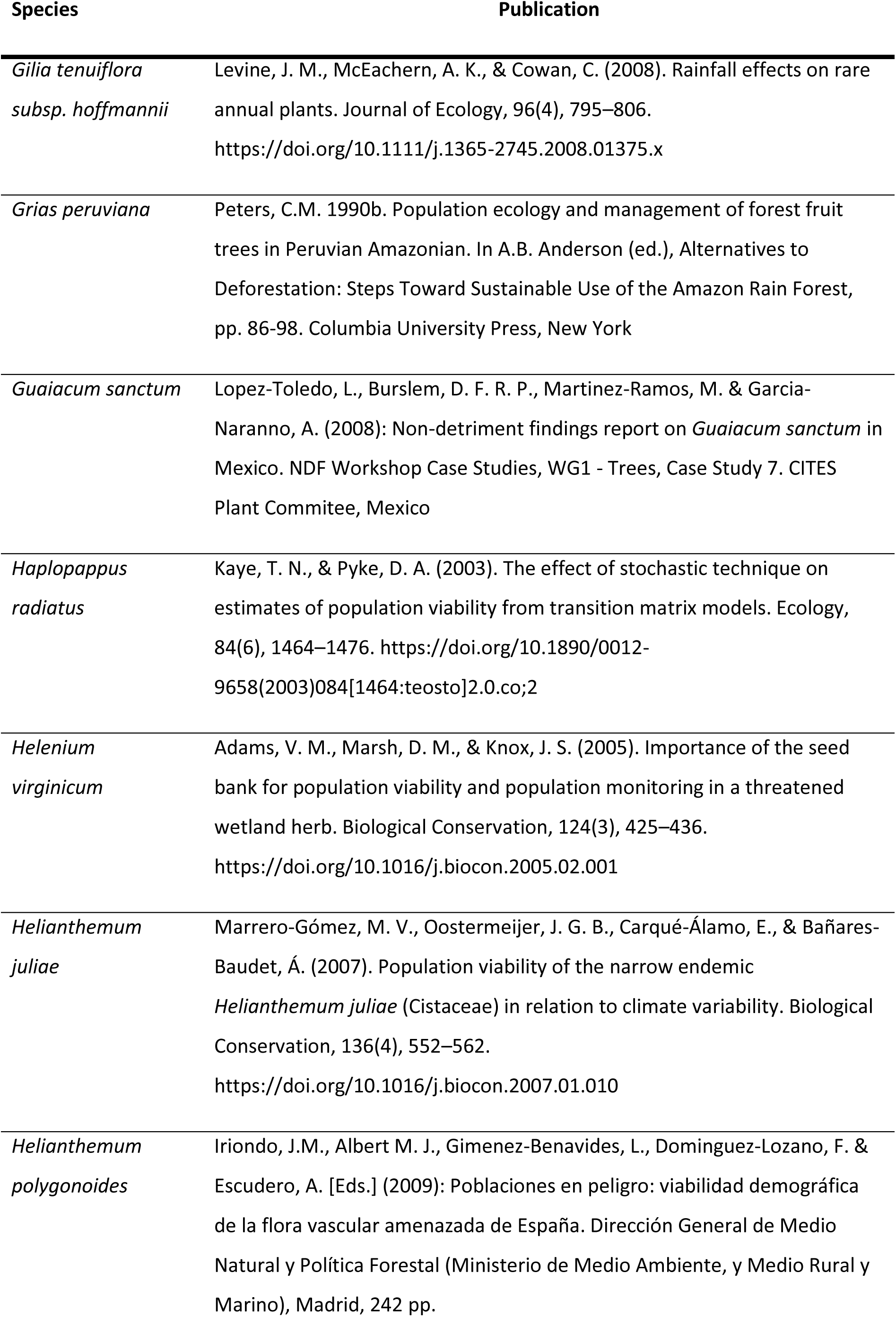
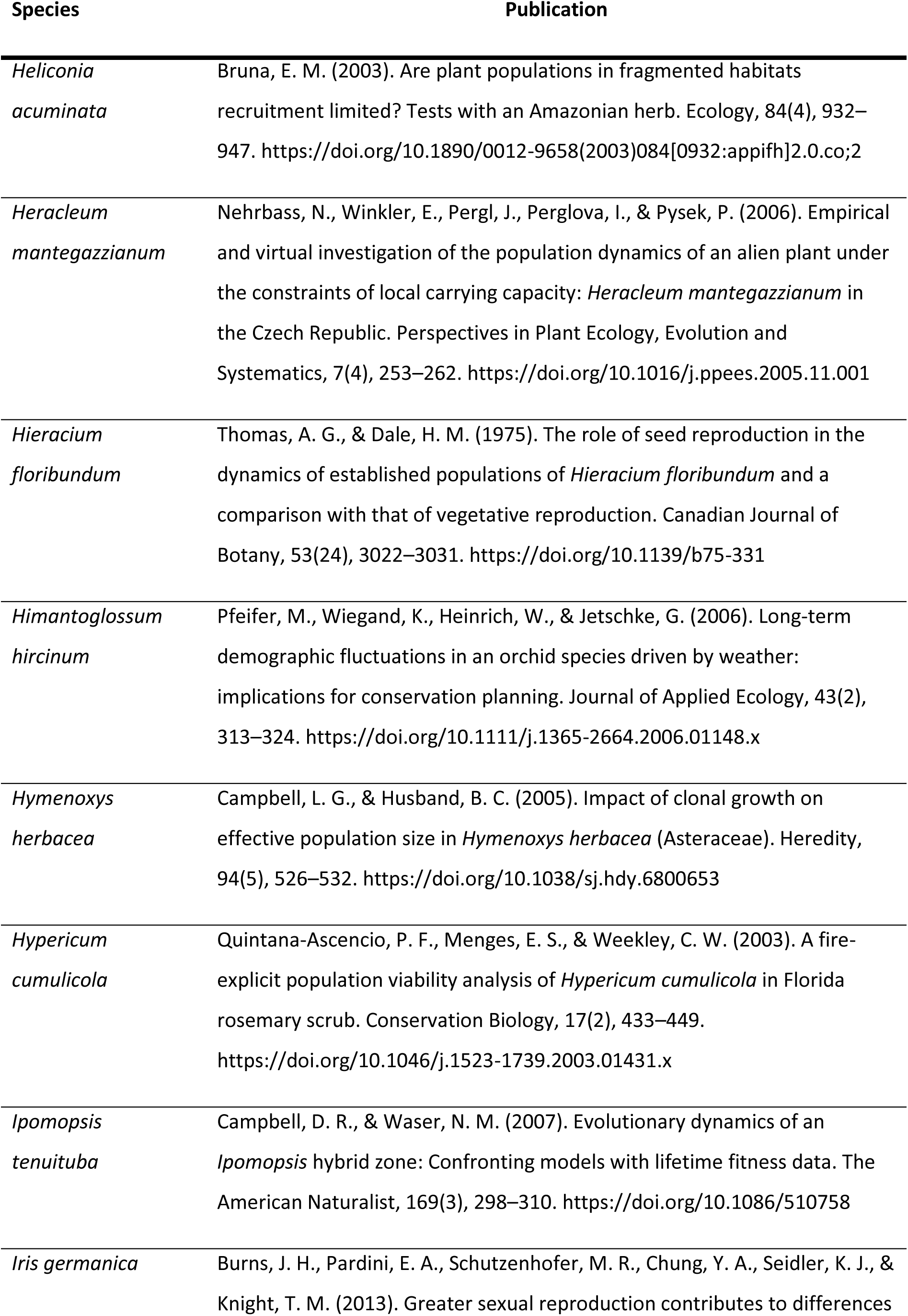
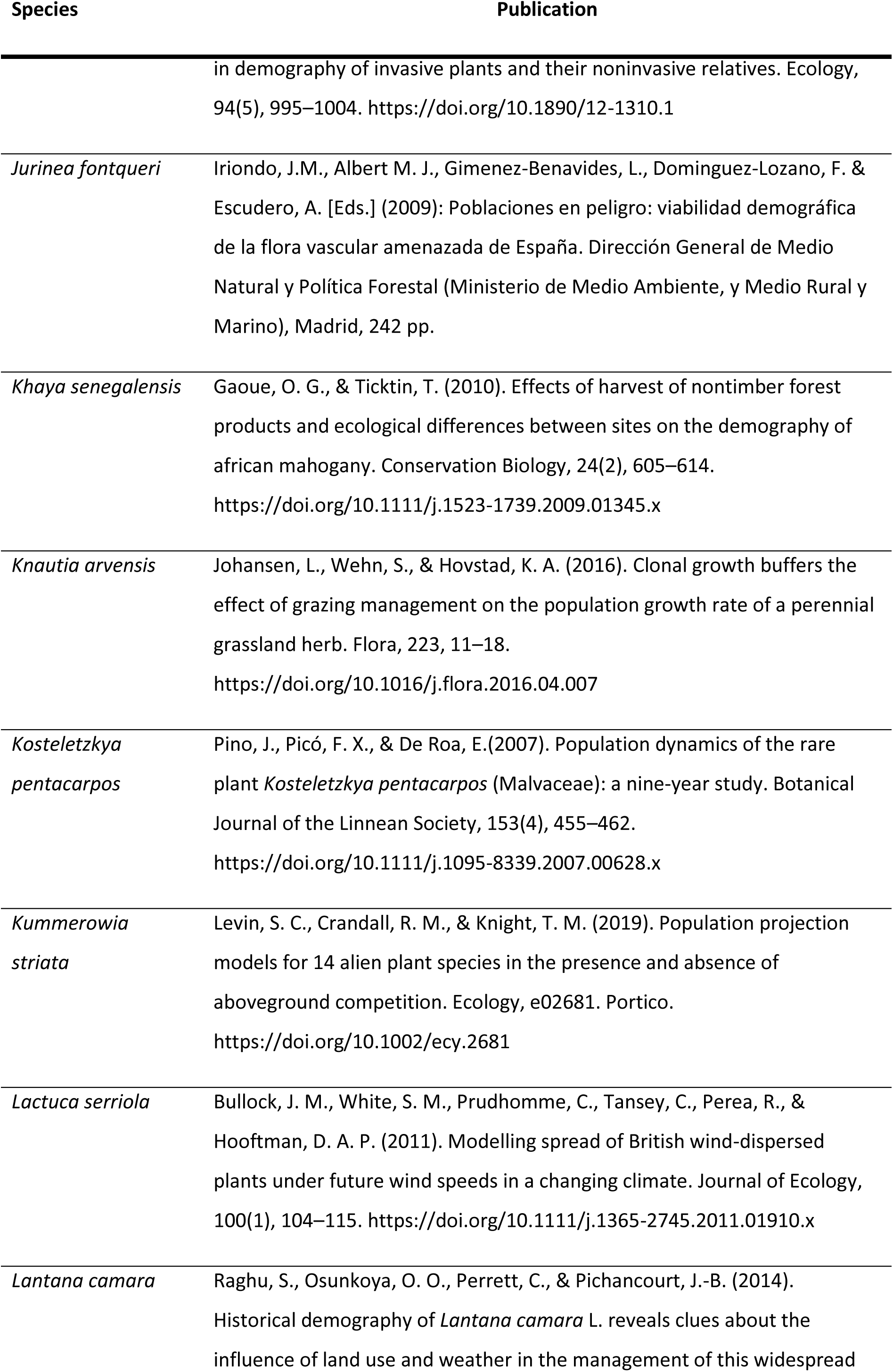
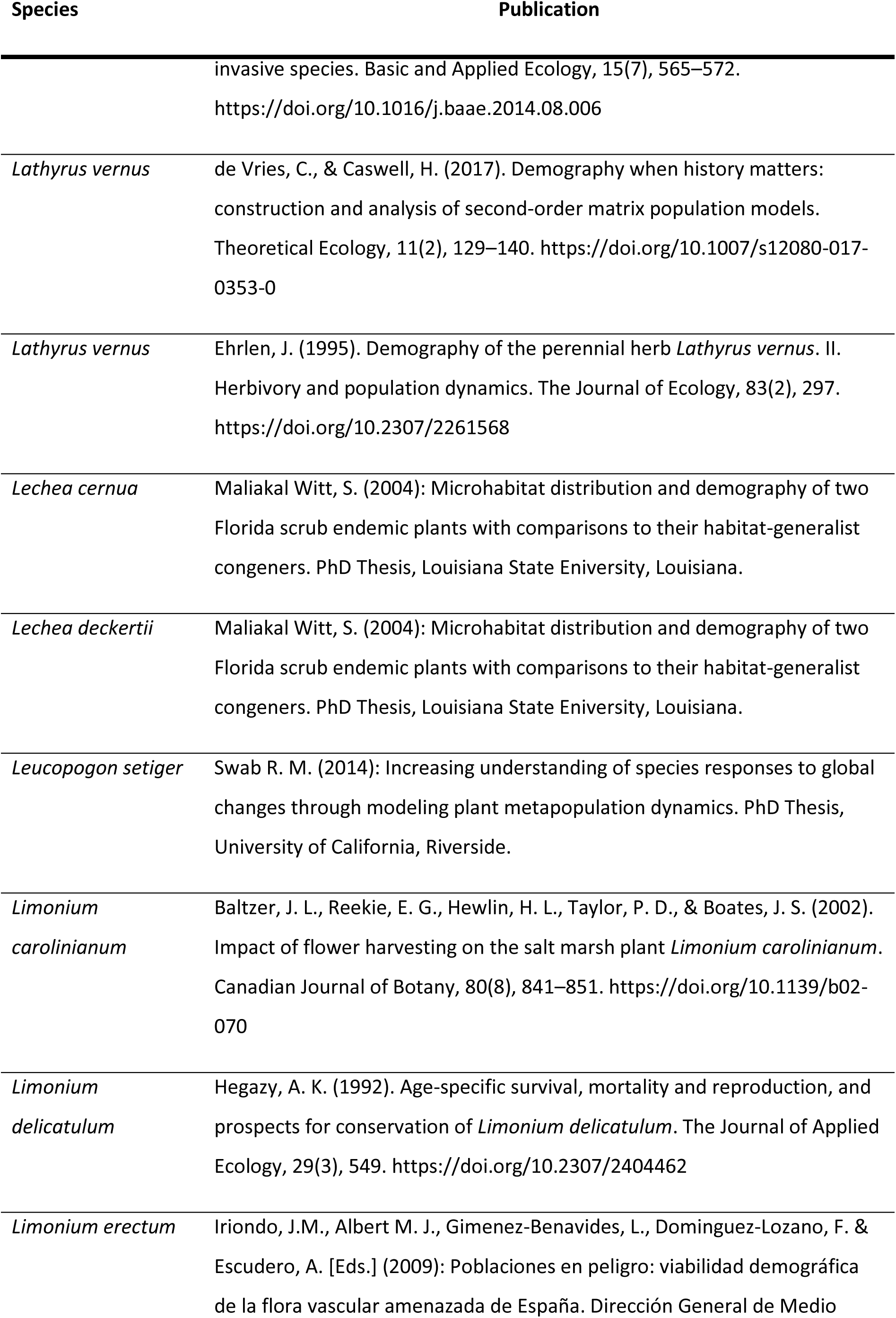
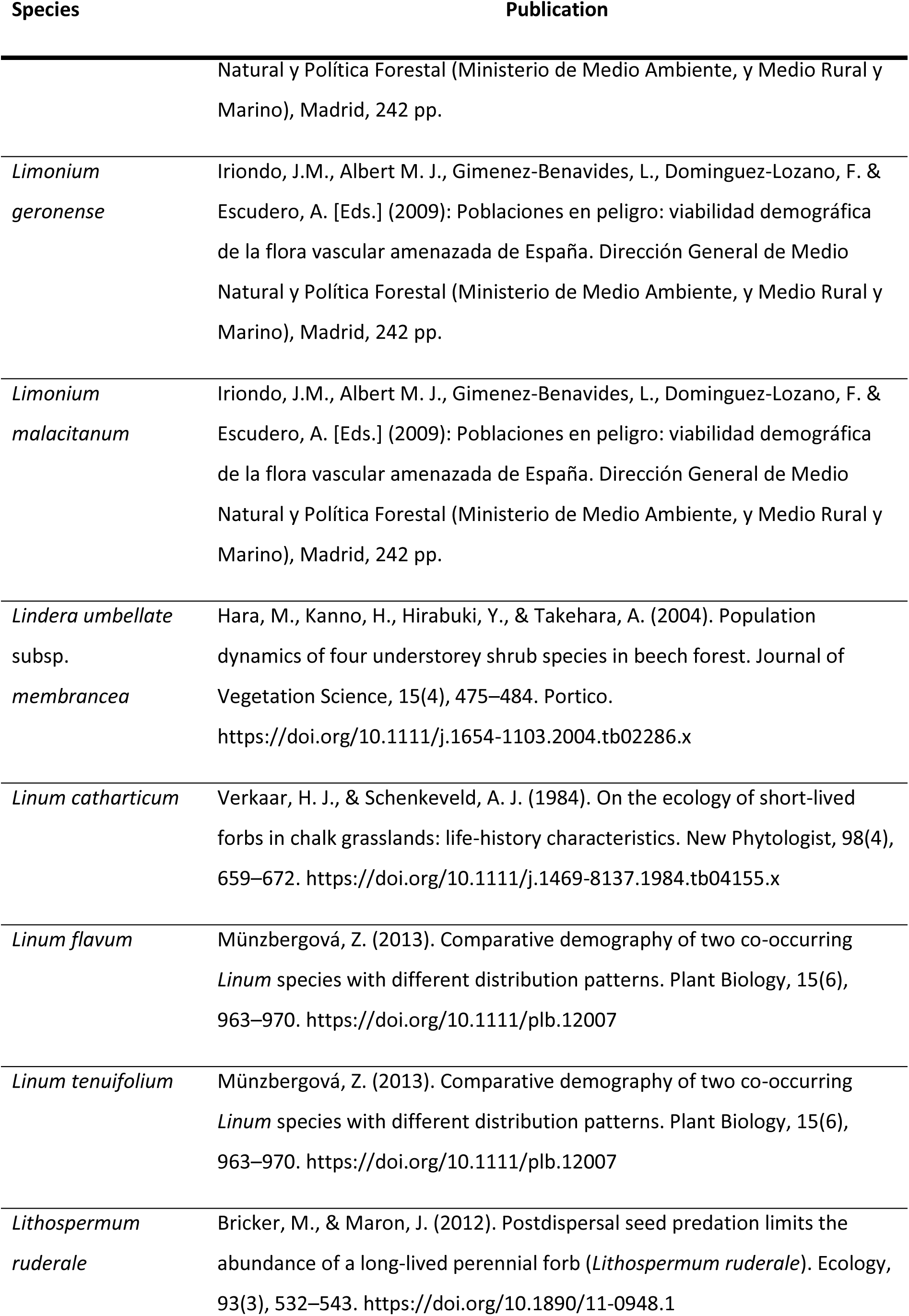
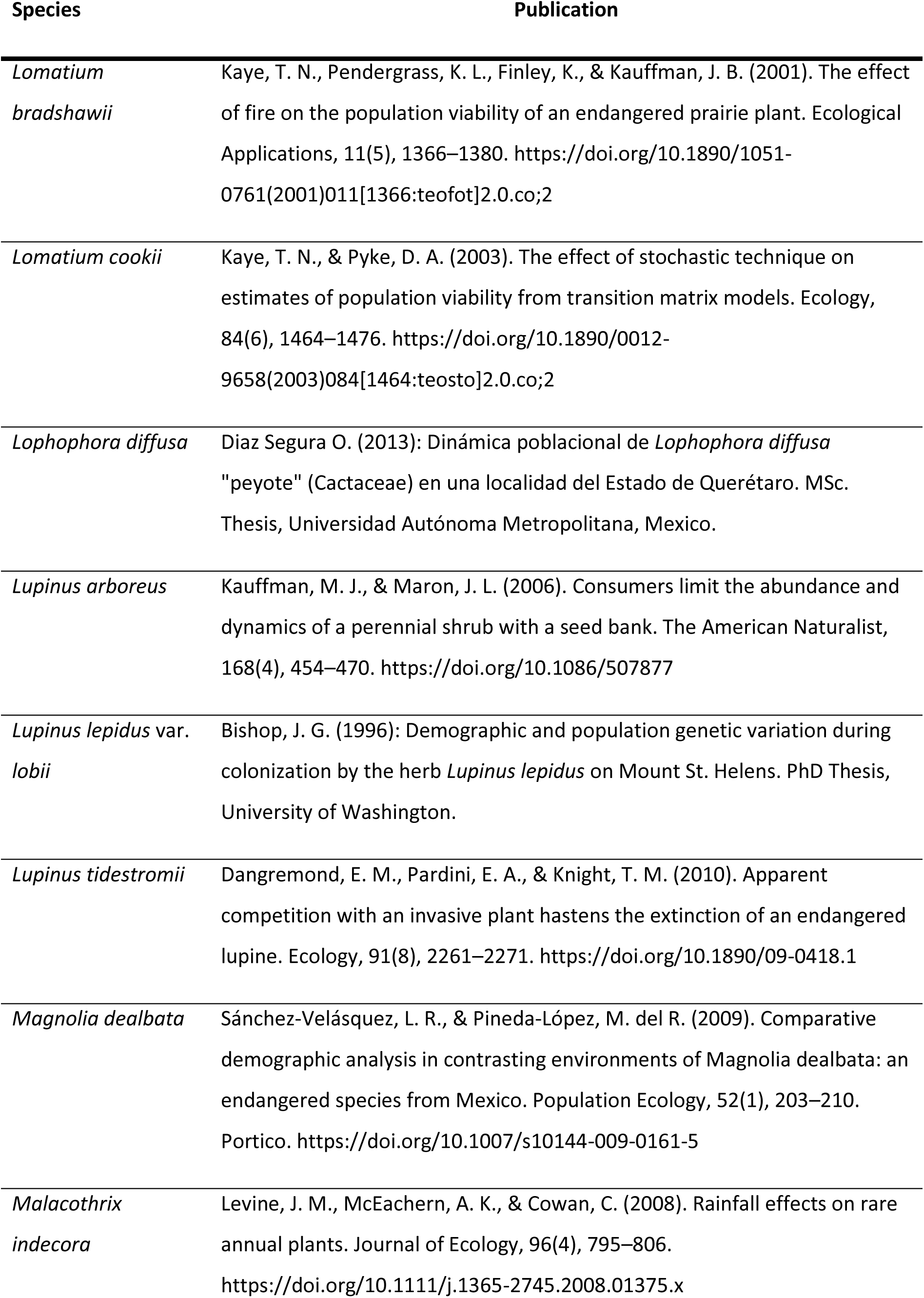
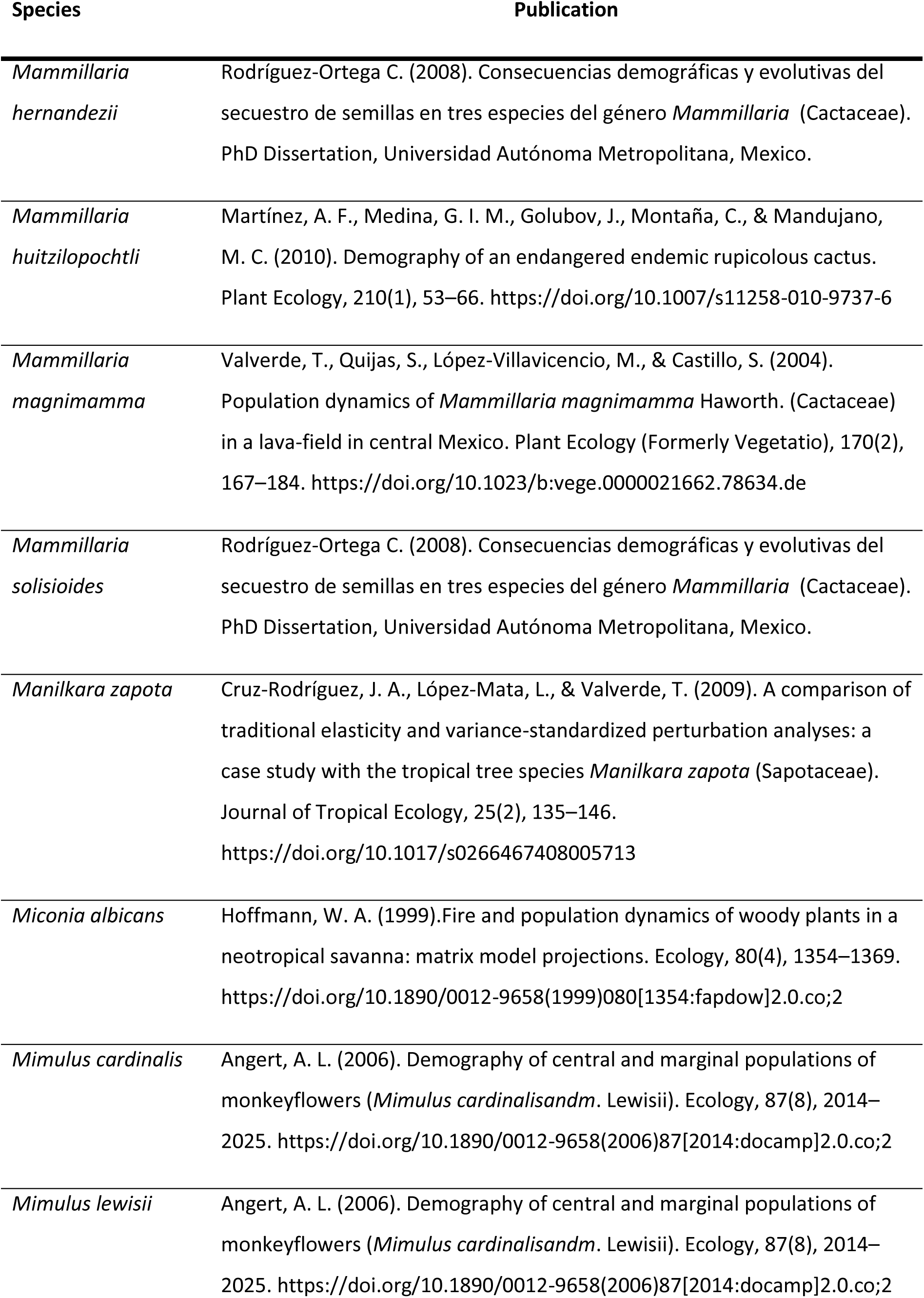
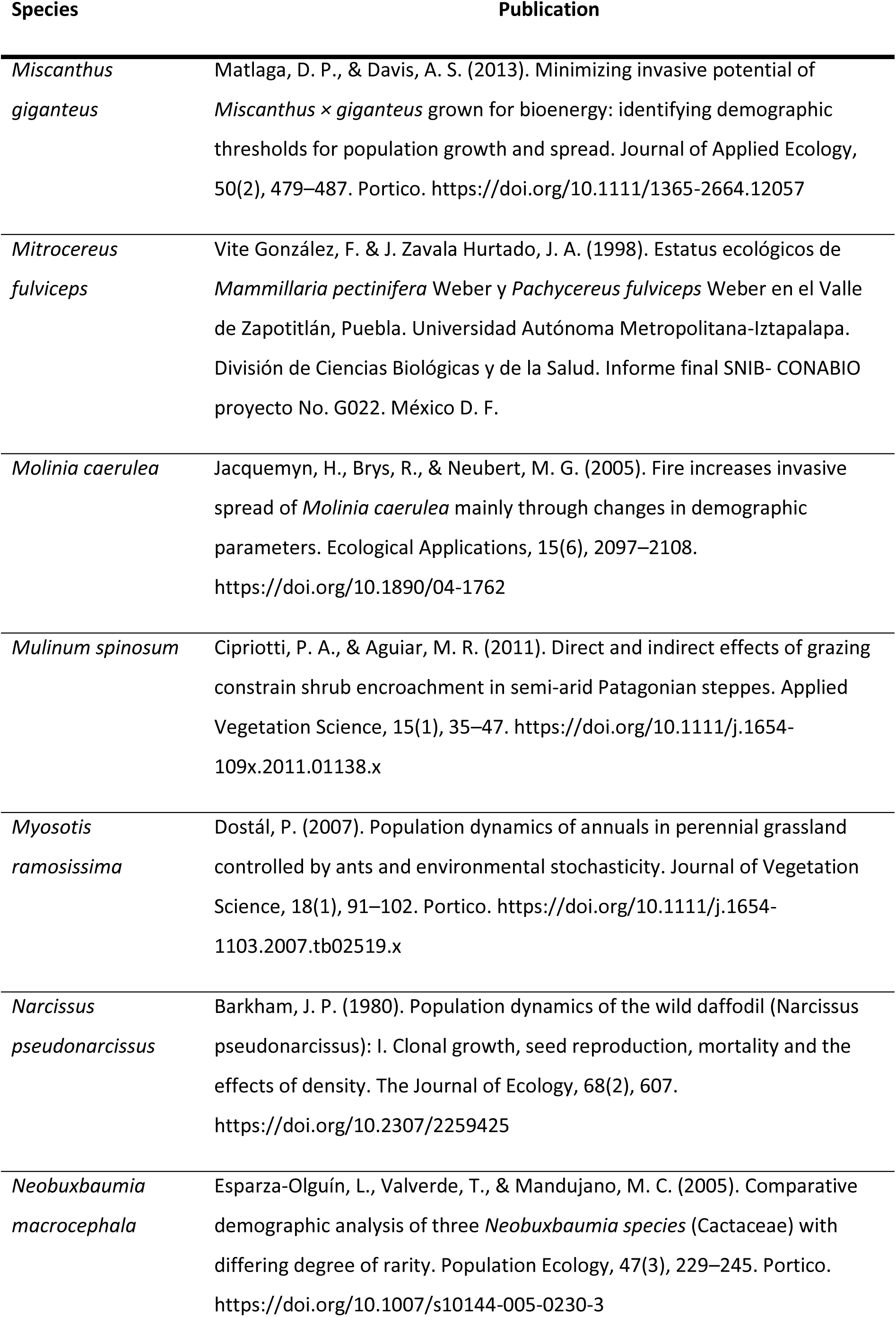
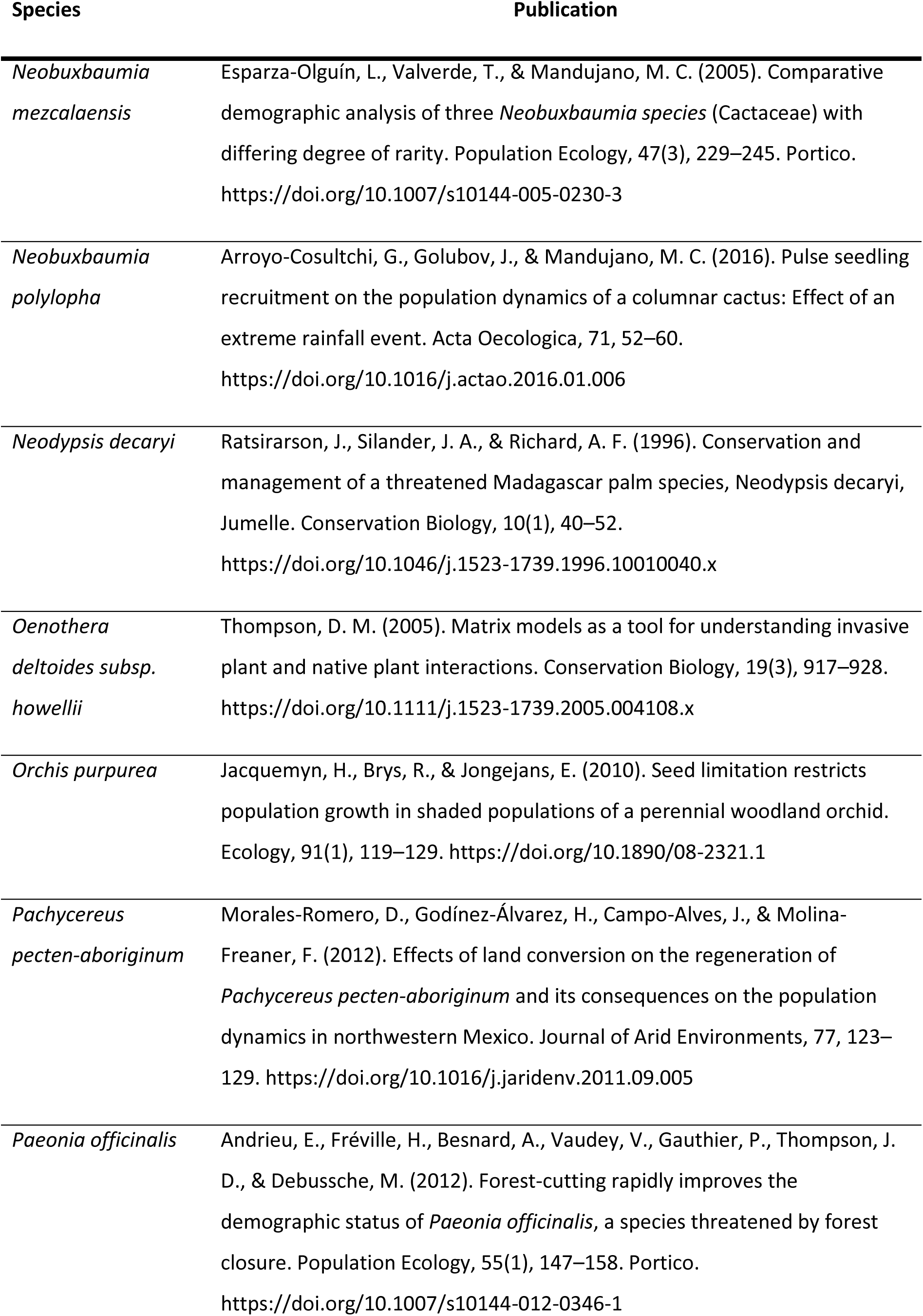
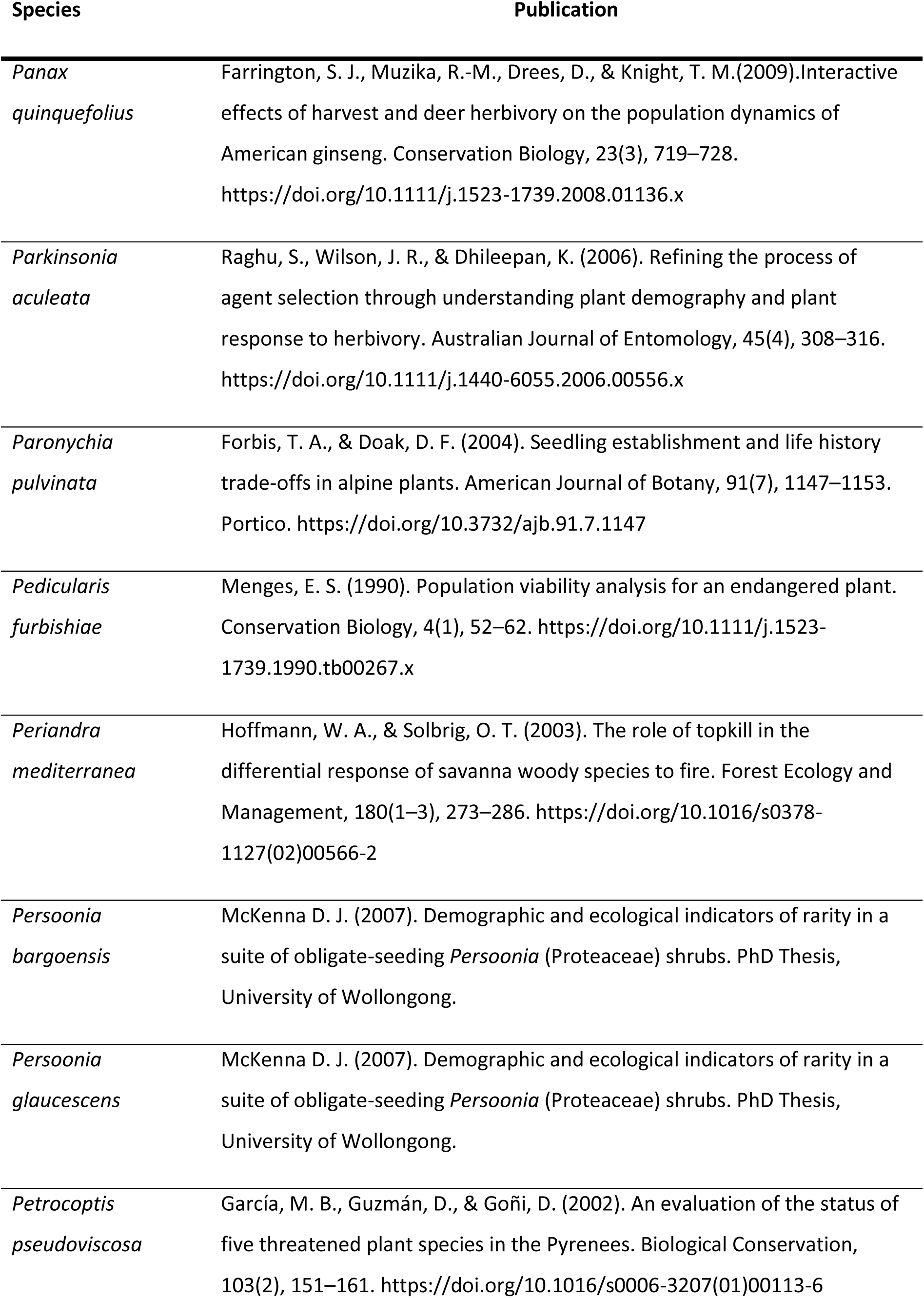
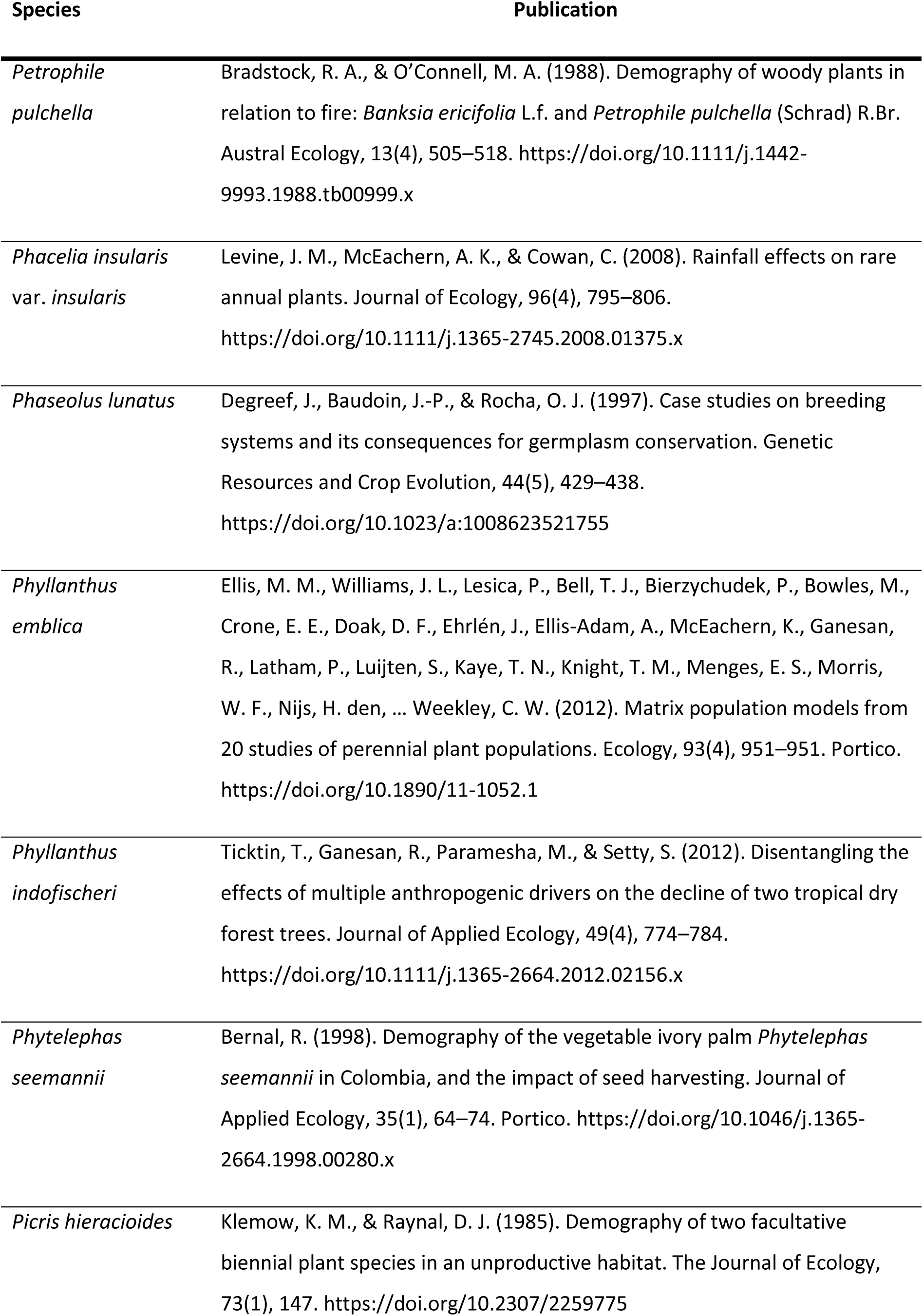
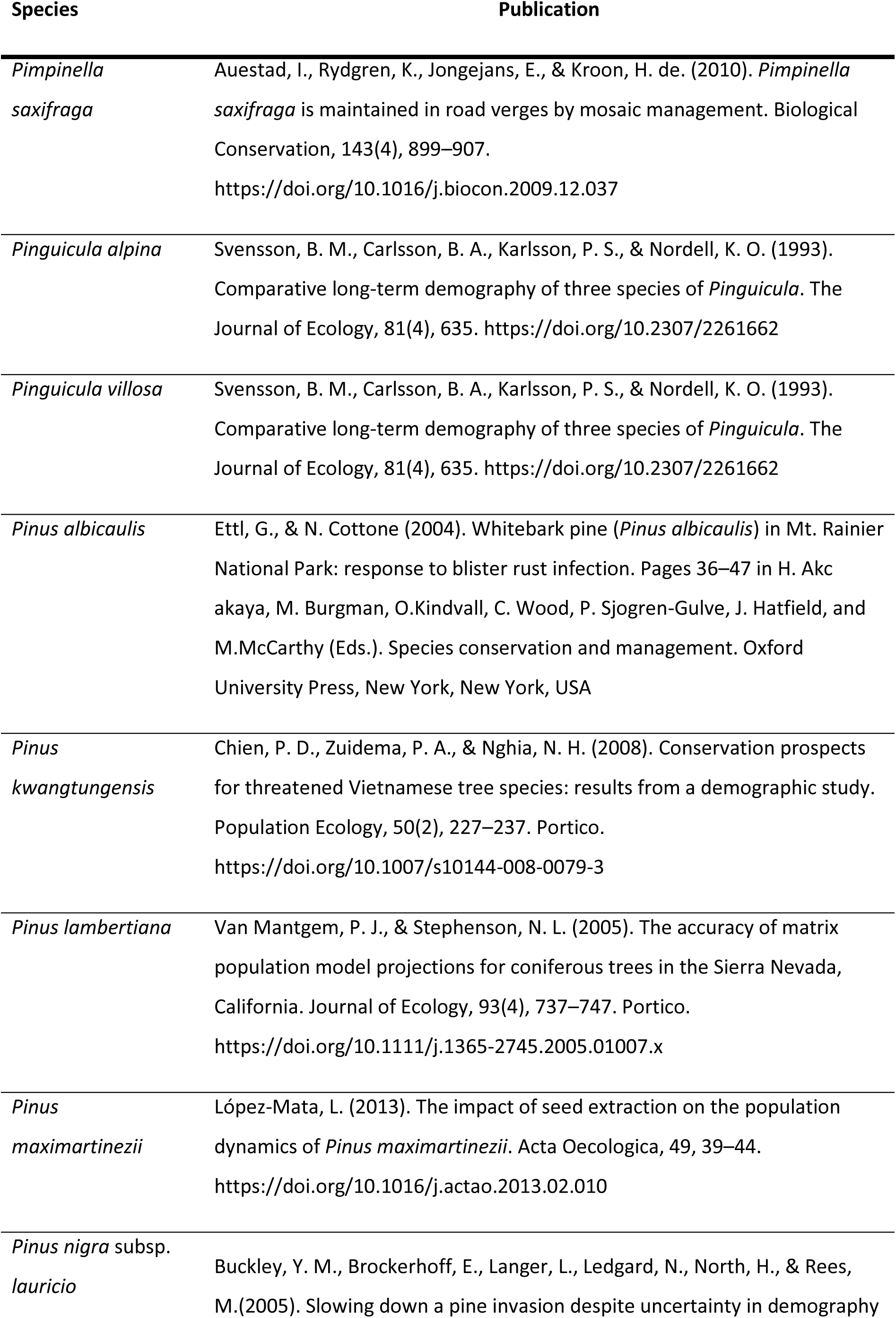
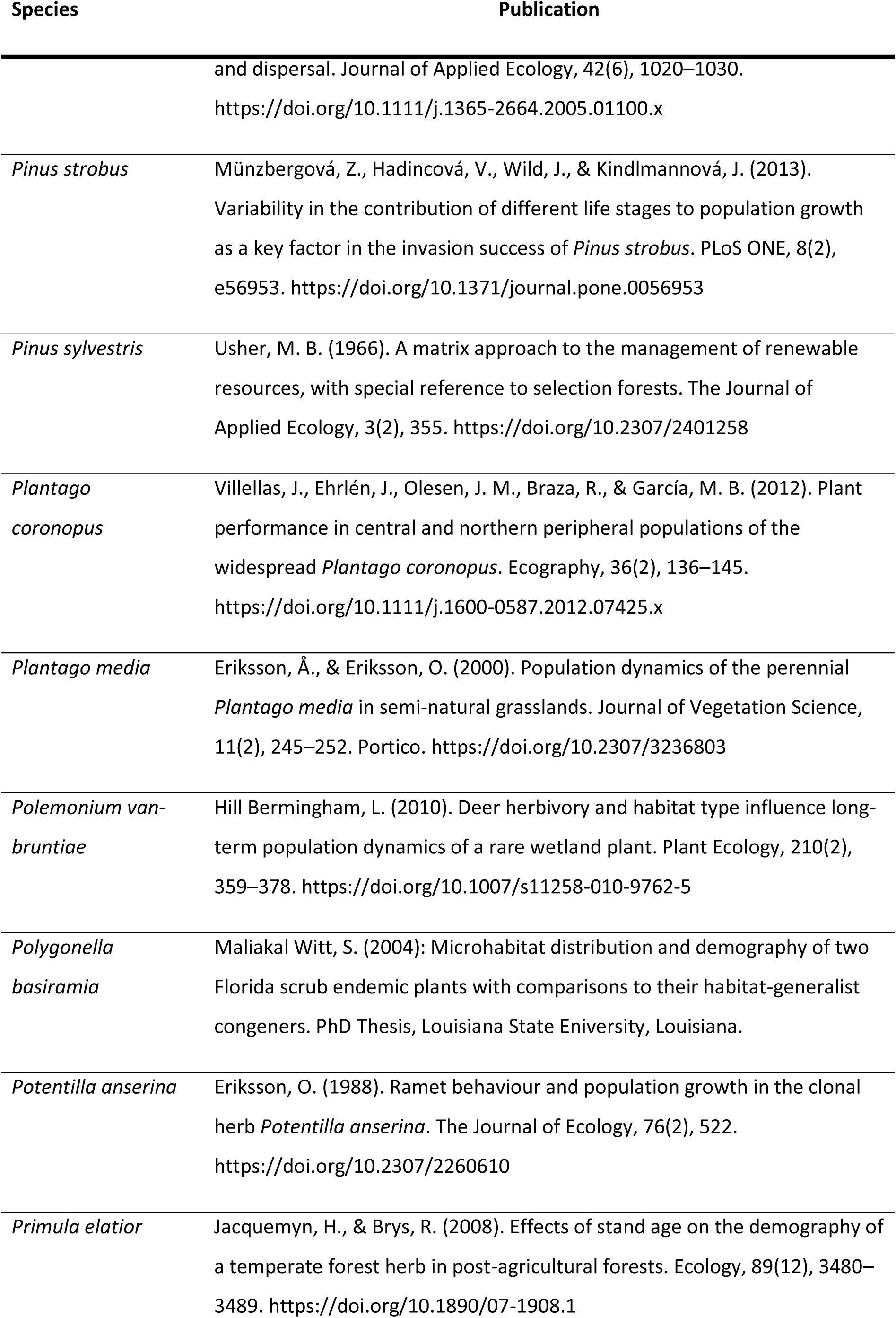
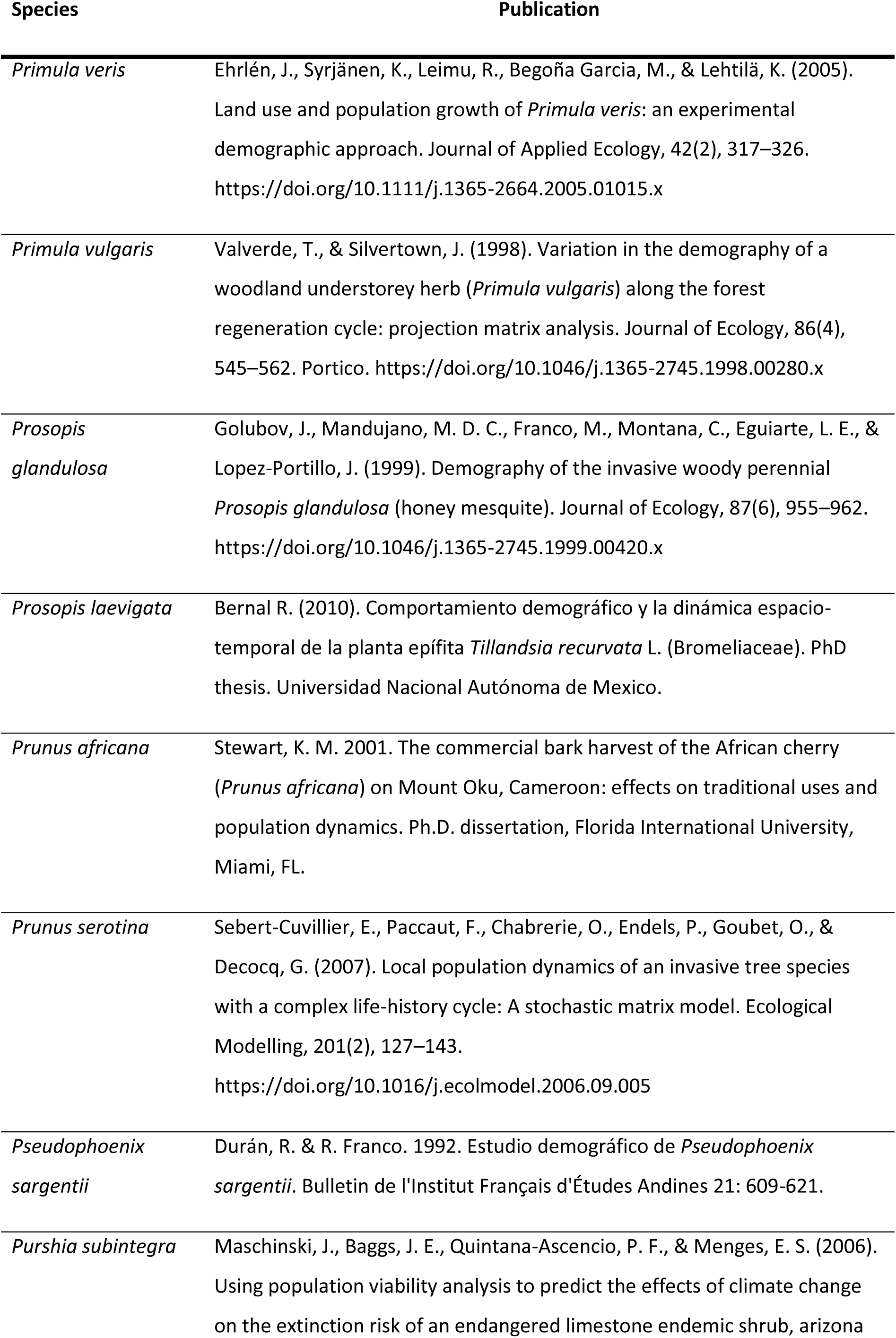
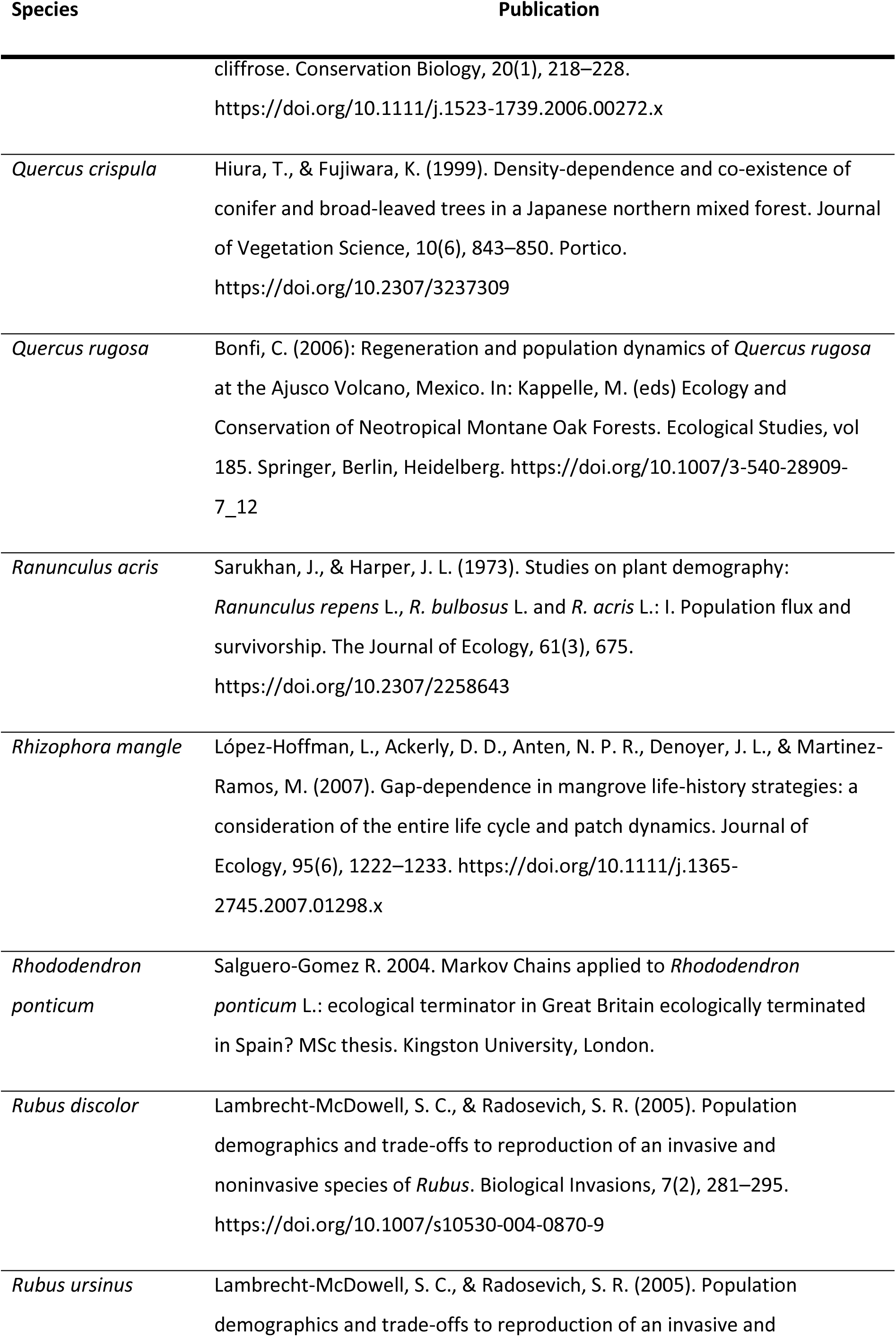
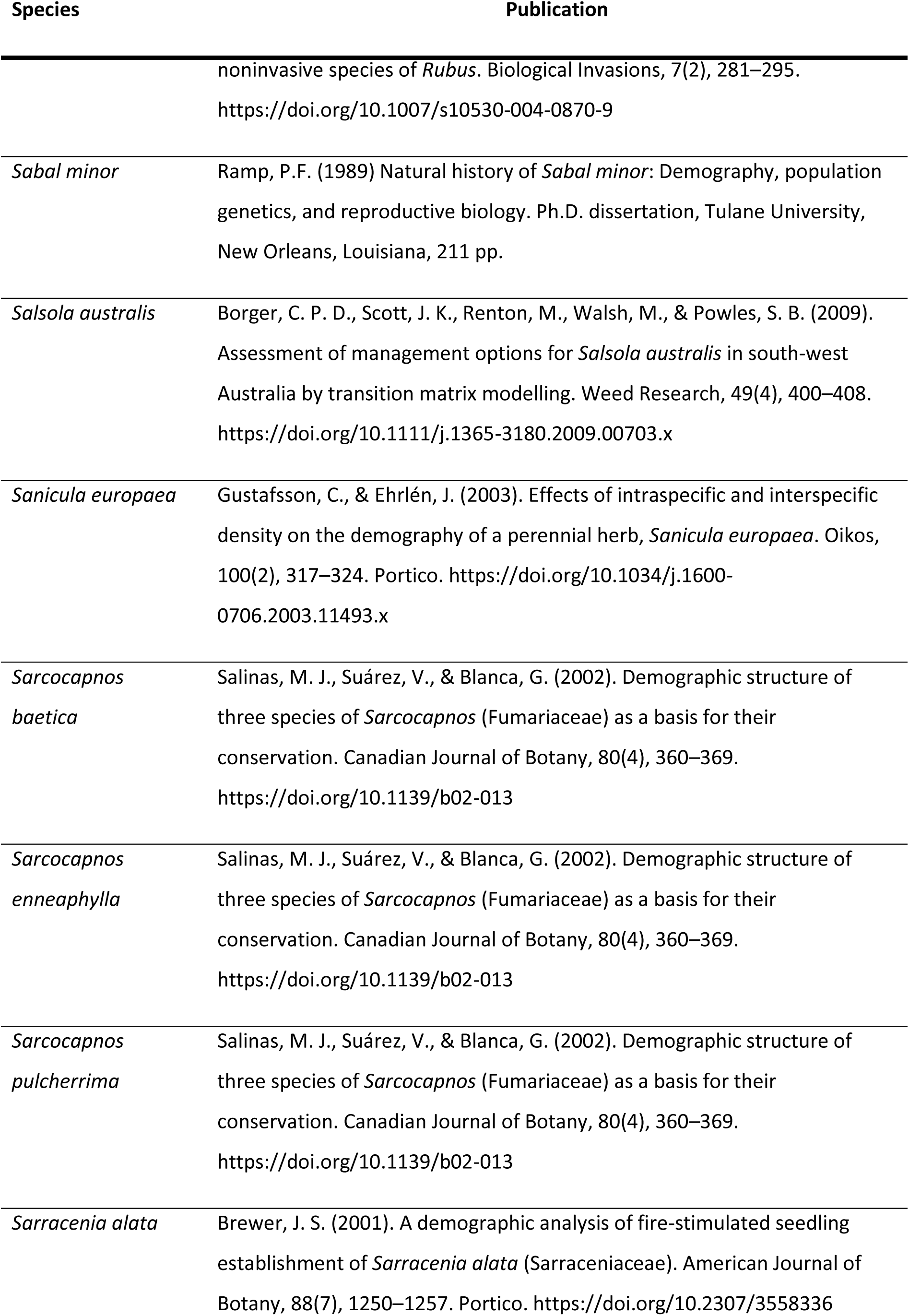
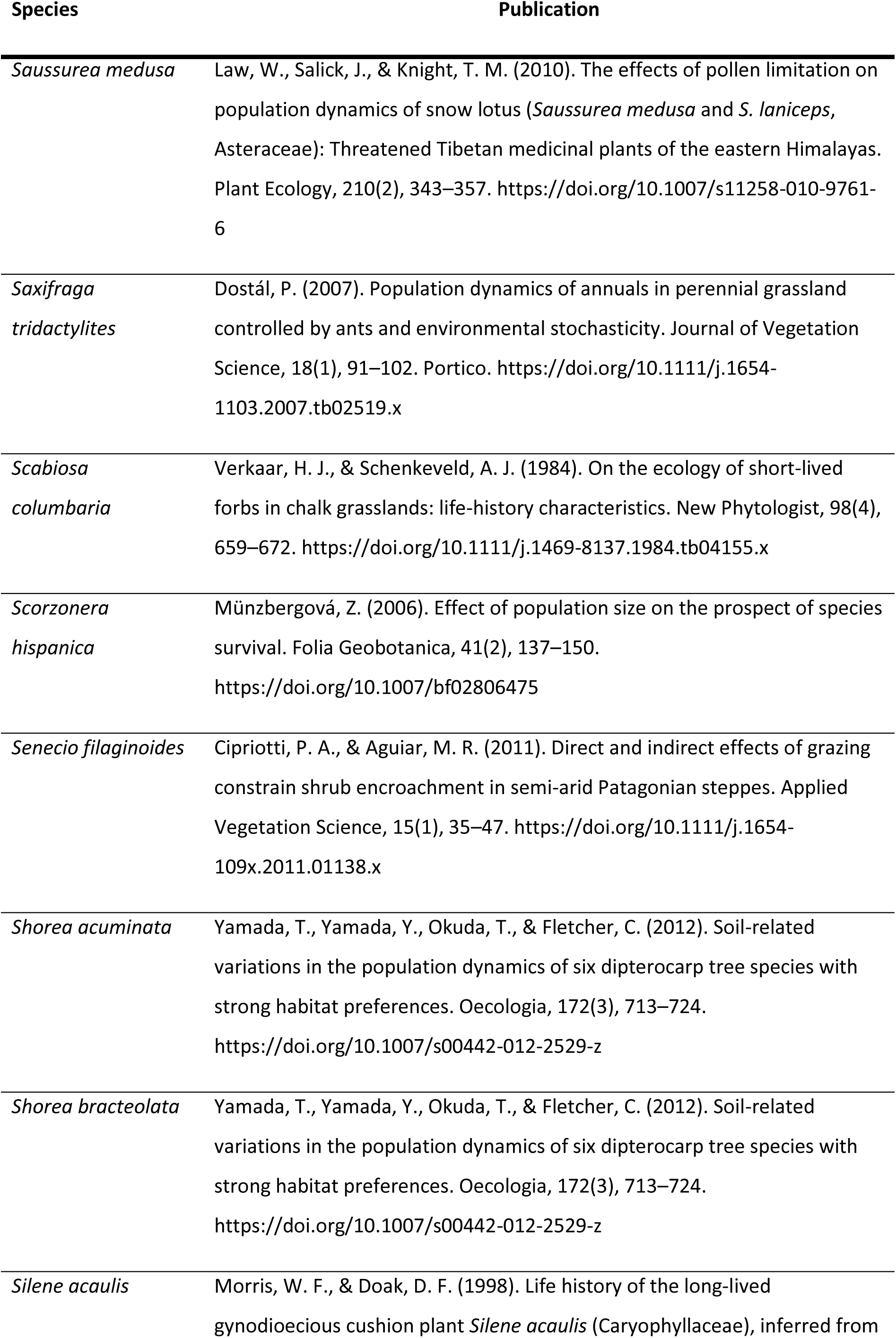
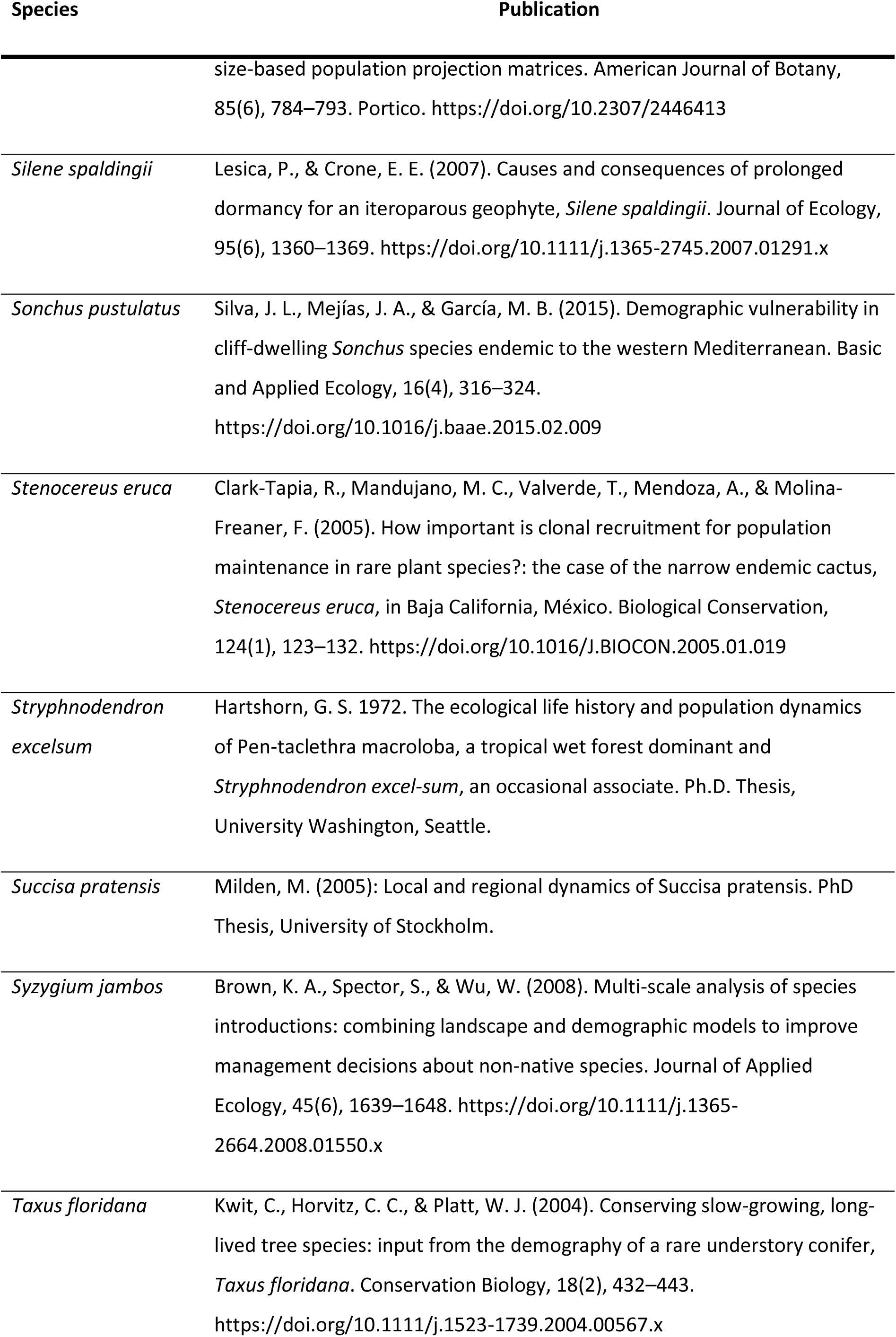
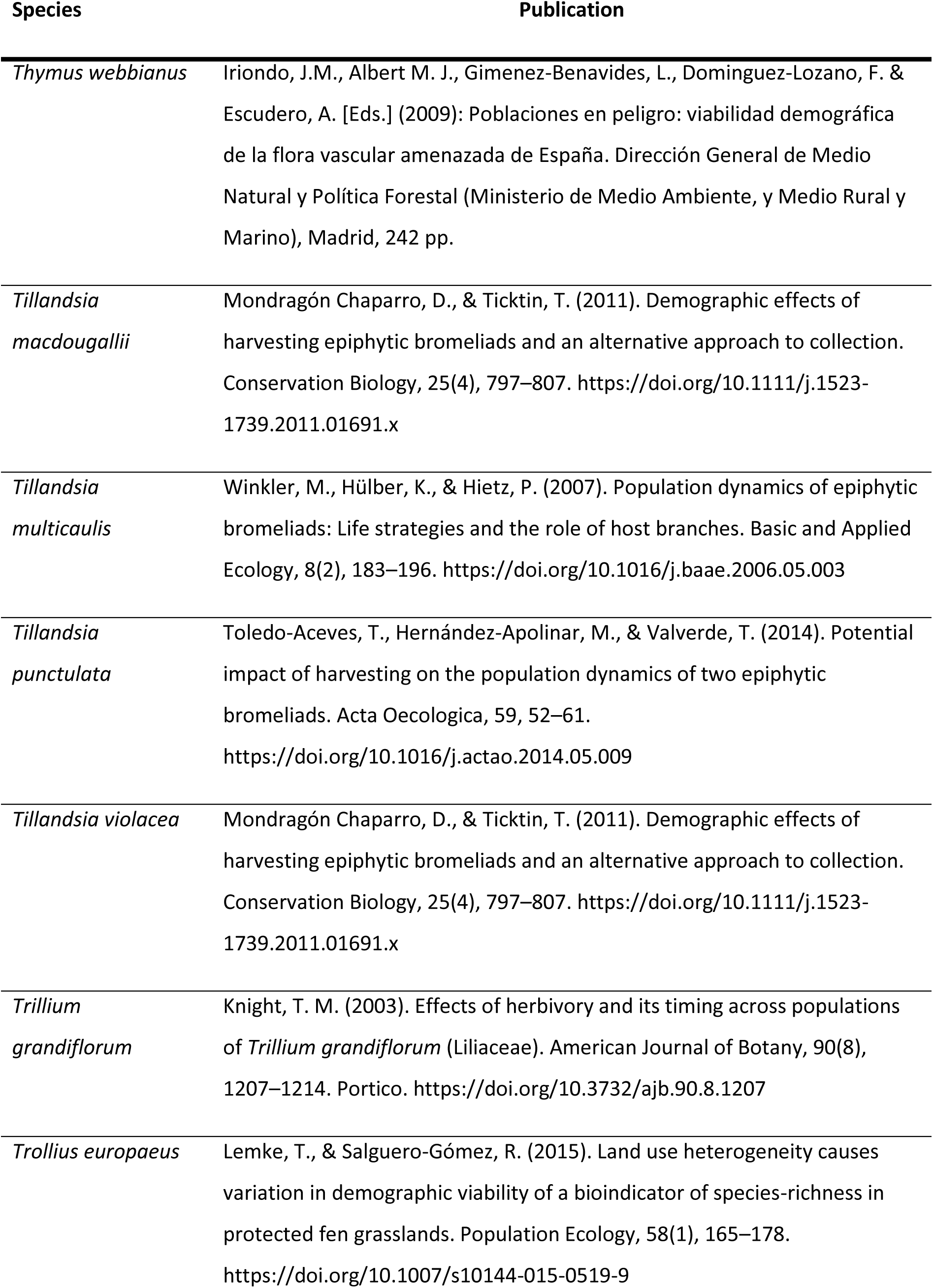
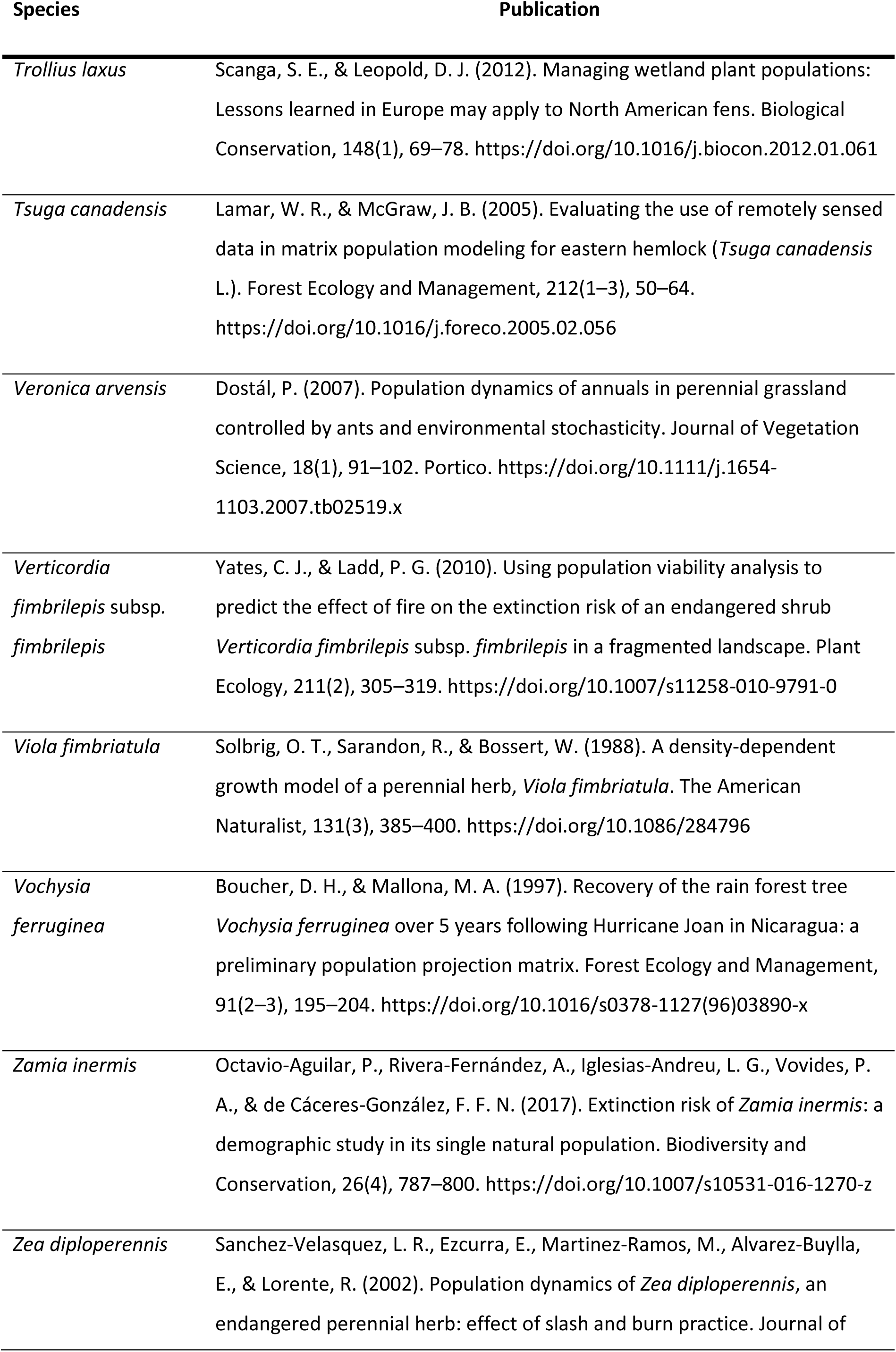
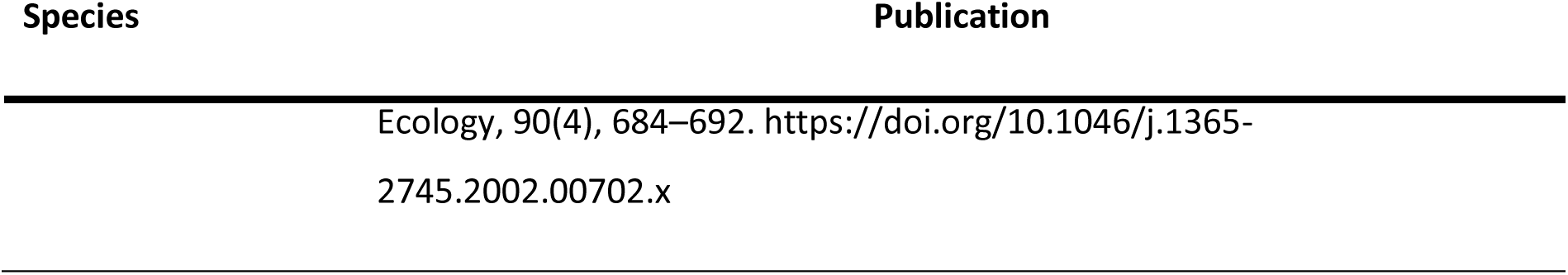
The final set of species used in this study and the original study that was the source of the MPMs.

**Table S2:**
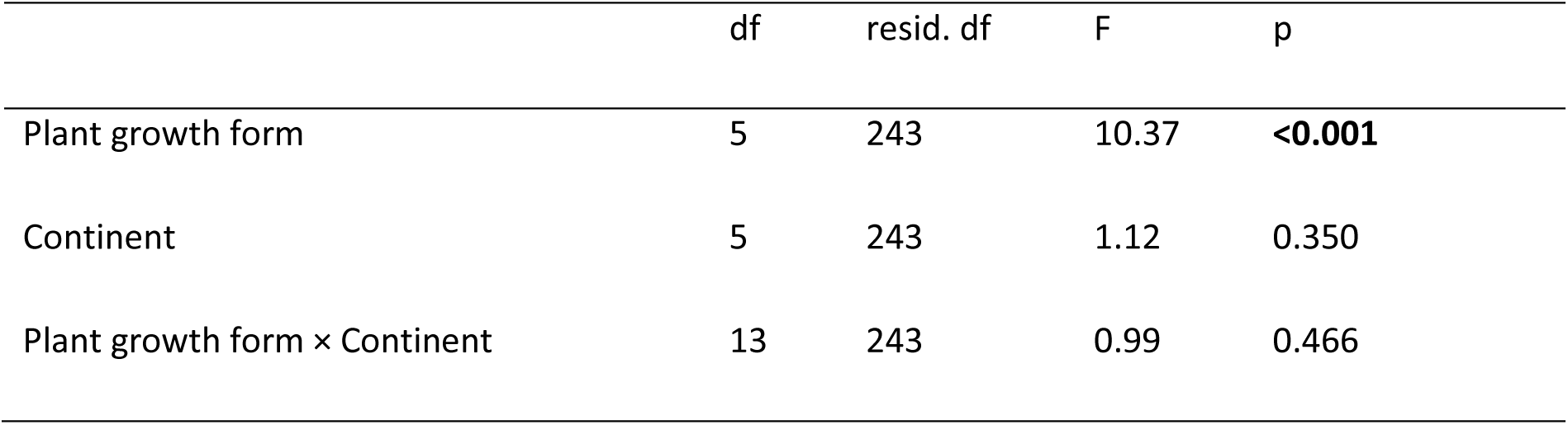
The effects of continent and plant growth form on vulnerability to seed harvesting. Results of a linear model with vulnerability to seed harvesting (log-transformed) as a response variable and plant growth form, continent, and their interaction as explanatory variables. We report results of a simple linear model because a generalized least square model with phylogenetic correction failed due to singular fit. The terms were fitted sequentially. Significant values are in bold. Adjusted R^2^=0.15

|  | df | resid. df | F | p |
| --- | --- | --- | --- | --- |
| Plant growth form | 5 | 243 | 10.37 | **<0.001** |
| Continent | 5 | 243 | 1.12 | 0.350 |
| Plant growth form × Continent | 13 | 243 | 0.99 | 0.466 |

**Table S3:** Formulation of the life history traits used to explain species vulnerability to seed harvesting in 280 vascular plant species. *λ* is the population growth rate, which corresponds to the dominant eigenvalue of the matrix ***A***; l_x_ and m_x_ are stage-specific survival and fertility schedules, ***C*** is the submatrix describing clonal reproduction, m is the dimension of the matrix ***C***, ***w*** is the stable stage distribution of the matrix ***A***, with j column entries of the matrix population model.

| Life history trait | Biological meaning | Formula |
| --- | --- | --- |
| Generation time $T$ | Number of years necessary for the individuals of a population to be fully replaced by new ones | $T = \frac{\log(\int_1^{\infty} l_x m_x dx)}{\log(\lambda)}$ |
| Degree of iteroparity $S$ | Spread of reproduction throughout the lifespan of the individual as quantified by Demetrius' entropy ($S$). High/low $S$ values correspond to iteroparous/semelparous populations | $S = -e^{-\log \lambda} l_x m_x \log(e^{-\log \lambda} l_x m_x)$ |
| Age at sexual maturity $L_\alpha$ | Number of years that it takes an average individual in the population to become sexually reproductive | $L_\alpha$ as described in Caswell 2001's equation 5.41 (35) |
| Seed bank residence | Mean amount of time individuals are expected to stay in the seedbank stage | As described in Caswell 2001's equation 5.36 (35) according to the fundamental matrix approach for the life cycle stage(s) that correspond to the seed bank stage(s) |
| Clonality $K$ | Per-capita clonal contributions weighted by the stable stage distribution of the MPM | $K = \sum_1^m \bar{c}_j \bar{w}_j$ |

**Table S4:**
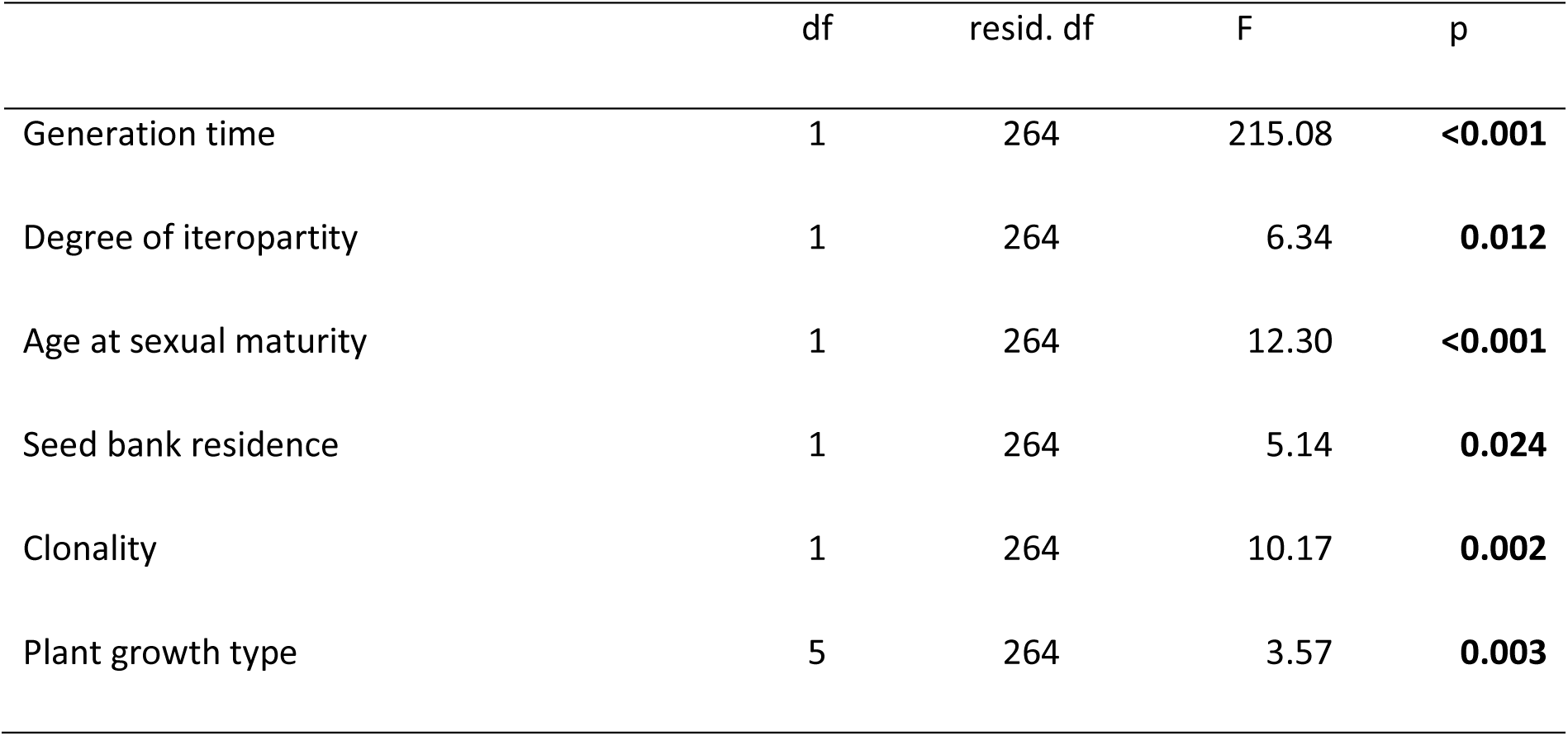
The effects of life history traits on vulnerability to seed harvesting, with significant values (*P*<0.05) in bold. Adjusted R^2^ = 0.61.

